# An empirical evaluation of simulated gastrointestinal digestion platforms for use in plant breeding, using common bean (*Phaseolus vulgaris* L.) as a model

**DOI:** 10.1101/2024.05.09.592089

**Authors:** Tayah M. Bolt, Margaret Riggs, Weiyi Sun, Li Tian, Paul Gepts, Antonia Palkovic, Travis Parker, Gail M. Bornhorst, Christine Diepenbrock

## Abstract

There remains a disconnect in plant breeding between increasing nutrient levels in crops at time of harvest and increasing bioaccessible levels of those nutrients during digestion. This study aims to develop and compare simulated digestion models for use in plant breeding and examine bioaccessible nutrient levels in common bean samples with differing seed coat coloration and patterning. The highest trait values (starch and protein hydrolysis, total phenolics, and antioxidant power) were observed from more dynamic digestion models, but even simple dynamic models showed higher trait values than a commonly used static digestion model. The use of these models provided insight on nutrient bioaccessibility; e.g., differences were observed during digestion between common bean genotypes for protein hydrolysis and between growing environments for both total phenolics and protein hydrolysis. Together, these results inform potential future pathways for applying simulated digestion models in plant breeding to improve bioaccessible nutrient levels in crops.

## 1 Introduction

Plant breeding is a critical tool for improving food quality, as the improvement is bred into the plant propagule (e.g., seed) without necessarily relying on further inputs or interventions later in the supply stream. However, if nutrient levels in the edible portion of the crop are assayed at all, they are typically solely measured at the time of harvest. This approach fails to account for the many physical and chemical changes that occur between harvest, processing, consumption, and nutrient absorption in the human body.^1^ Consequently, measures of nutrient levels at time of harvest are not necessarily representative of the levels of bioaccessible nutrients (i.e., nutrients that are available for the human body to absorb). Closing this disconnect is critical to maximize the improvements to human nutrition that are achievable and attained through plant breeding.^2,3^

Accurately measuring the bioaccessibility of a given nutrient is a complex process and a major reason why nutrient levels are typically only measured at time of harvest, if at all. To measure bioaccessibility, the dynamic digestive process and its impacts on the food matrix and subsequent release of nutrients from that matrix must be considered. Ideally, *in vivo* studies would be used to understand this complex process; however, these can be cost-prohibitive and require human test participants.^4^ Most commonly, simplistic static *in vitro* methods are used in lieu of *in vivo* methods. A limited number of dynamic models have been developed and include features such as pH regulation, food flow (including gastric emptying), peristaltic (wavelike) contractions of the stomach wall, and real-time injection of digestive fluids which help to better simulate the complexities of the human digestion process.^5^ The Human Gastric Simulator (HGS) used in this study is one such model that incorporates mechanical peristaltic contractions, as well as dynamic secretion of simulated gastric fluids and gastric emptying of the meal.^5–8^ Additionally, a multi-channel Peristaltic Simulator (PS) was developed to allow for higher throughput compared to the HGS, and allows for running up to 12 samples simultaneously.^9^ In that study, the PS was found to produce degree of occlusion of the peristaltic wave, pressure applied to a bolus, and fluid velocity within physiologically relevant ranges. The high-throughput PS model could render the use of dynamic digestion methods more feasible for routine use in breeding programs, which are typically producing (in replicated field trials) and analyzing hundreds to thousands of samples per year.

Common bean (*Phaseolus vulgaris* L.) was used as an experimental system herein. Common bean is the most important grain legume for direct human consumption,^10^ and includes many commercially relevant market classes (e.g., kidney, pinto, black, and navy, in the U.S. alone) that are under active improvement by plant breeders. Dry beans (i.e., common bean seeds harvested from mature pods) are a primary source of protein in parts of Central America, South America, and East Africa and also provide dietary fiber and certain micronutrients.^11–14^ Protein and starch hydrolysis are of relevance for human nutrition both in whole beans and when using beans as ingredients for the nutritional improvement of food products.^15^ Increased protein hydrolysis and digestibility particularly of phaseolin, a seed storage globulin that comprises approximately 30-50% of total protein,^16–18^ has been of interest in breeding to increase levels of free amino acids and bioactive peptides and potentially mitigate tradeoffs between total protein and yield.^19,20^ Protein digestibility in common bean has been assayed *in vivo* in animal models and *in vitro* using static, but not dynamic, digestion methods.^21,22^ Decreased rate and extent of starch hydrolysis could be helpful for glycemic control among other health benefits, as starch that is resistant to digestive enzymes before and during small-intestinal digestion (rather is fermented in the colon or excreted), classified as resistant starch, is a type of dietary fiber.^23,24^ Gallegos-Infante et al. found that pasta made with added common bean flour had both higher protein levels and slower starch hydrolysis rates than the pure semolina control.^25^

Seed coat coloration and patterning are often used by consumers to identify their desired market class and could affect the appearance of the final food product.^26,27^ As consumers and growers often exhibit strong preferences for dry beans with specific seed coat coloration/patterning, a vast diversity of seed types can be observed in common bean and related grain legumes.^28^ This diversity provided an opportunity to assess bioaccessible nutrient levels through simulated gastrointestinal digestion in samples of common bean with contrasting seed coat coloration/patterning. This coloration and patterning are primarily underlain by phenolic compounds, which are among the largest classes of metabolites in plants. In common bean among other grain legumes, phenolics are primarily localized in the seed coat, and phenolic acids and flavonoids (including proanthocyanidins) are among the most abundant subclasses.^29–33^ While phenolics are potential contributors to the nutritional impacts of beans, the health benefits and antioxidant potential conferred by dietary phenolics are highly dependent on their absorption and metabolism throughout the digestive process.^11,34^ For example, in black beans it was found that some phenolic compounds inhibit iron uptake while others promote iron uptake.^35–37^ Phenolics can also interact with both starch and proteins,^38–41^ with multi-faceted impacts on nutritional outcomes and compositional traits.^42–46^ Bioaccessibility of phenolics in strawberry achenes (which are enriched in phenolics compared to the remainder of the fruit) and in Northern highbush blueberries have also been explored as breeding targets;^47,48^ however, those studies conducted *in vitro* digestion using shaking or oscillation without the use of a dynamic model that could more accurately replicate the conditions in the human gastrointestinal tract.^5^

Lastly, environmental effects are majorly important when considering plant breeding approaches and applications. For example, in common bean, it has been shown that the environment can majorly affect both seed coat coloration/patterning and seed zinc and iron concentrations.^49,50^ The main and interaction effects of genotype and environment on bioaccessible nutrient levels through simulated gastrointestinal digestion are yet to be characterized in several populations and germplasm pools of importance for common bean breeding and production.

This study aims to: 1) develop and compare dynamic digestive models for use in plant breeding; and 2) examine bioaccessible nutrient levels (for protein, starch, phenolics, and antioxidant power) in common beans with differing seed coat coloration/patterning. For our first aim, we compared four digestive platforms: a shaking water bath, the PS, the HGS with dynamic gastric secretions and emptying (hereafter referenced as the ‘full HGS’ method), and a simplified HGS method. For our second aim, two recently released common bean varieties (UC Southwest Red and UC Southwest Gold ^51,52^) that exhibit variability in seed coat patterning across field environments within California ^49^ were analyzed using the PS and simplified HGS platforms. These sample types (representing genotype-environment combinations) were used to examine gastrointestinal outcomes from beans with varying pigmentation hue (perceived coloration; red vs. gold) and percentage of pigmented seed coat area. Combined, these two aims allowed the outfitting and testing of dynamic digestive models for use in plant breeding while also gaining insight into nutrient bioaccessibility in common bean accessions that differ in seed coat coloration/patterning.

## 2 Materials and Methods

### 2.1 Overall Experimental Design

This experiment encompasses two principal aims: to compare differing digestion platforms, as seen in Figure 1, and to compare common bean varieties that exhibit differential extent of pigmented seed coat areas. The digestive platforms utilized in the first aim included a shaking water bath (static method), the PS, the full HGS, and a simplified HGS method. The second aim utilized two recently released common bean varieties that each exhibited differential extent of pigmented seed coat area across field environments within California that were analyzed using the PS and simplified HGS platforms, as these allowed for testing of genotype-environment combinations with limited available sample mass. For each digestion, samples were taken at a minimum of five time points for analysis and comparison. Each digestion platform was performed in triplicate from raw beans through to the small intestinal phase. All chemical sources can be found in Table S1.

**Figure 1.**
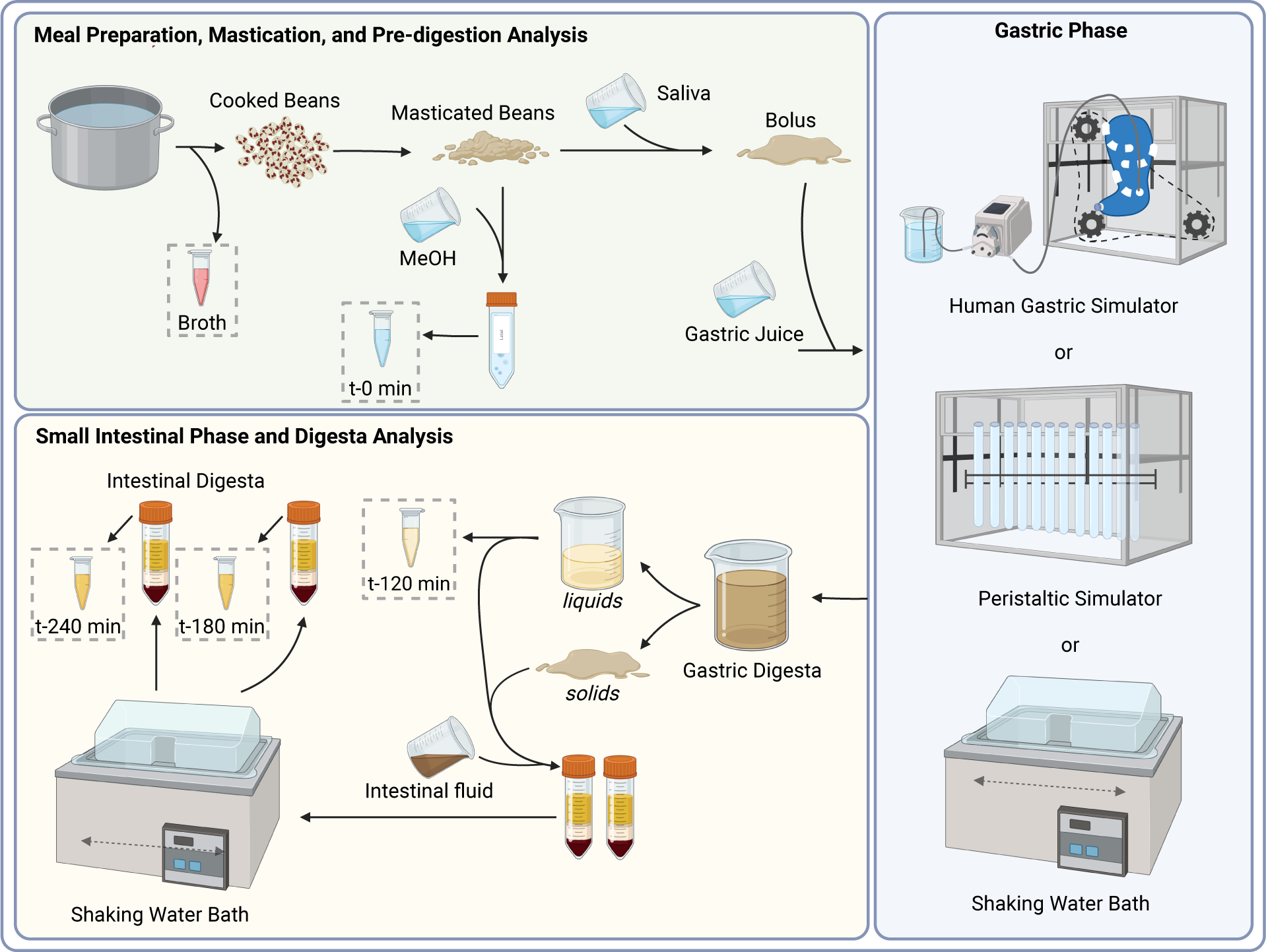
Digestion method workflow. Small tubes labeled Broth, t-0 min, t-120 min, t-180 min, and t-240 min (demarcated using dashed boxes) represent the liquid fraction of the samples which were utilized for further analysis. Abbreviations: MeOH, methanol; t-, time point.

### 2.2 Sample Selection

Two recently released common bean varieties (UC Southwest Red and UC Southwest Gold ^51,52^) that tend to exhibit variability in the extent of pigmented seed coat area across field environments within California ^49^ were used herein. UC Southwest Gold and Red are common bean lines descended from a cross between ‘Zuni Gold’ and UCD 9634 pink bean (and specifically, are the F_7_ bulk-harvested progeny of a single F_3_ progenitor and an F_9_ family from a single F_5_ progenitor, respectively ^51,52^). UC Southwest Gold seed that was produced in San Gregorio, California, in the Summer 2022 growing season was used for the four-way digestion method comparison (aim 1), as a large amount (more than the 900 g needed for three experimental runs on all four digestion methods) of that sample type was available for testing across the different simulated digestion methods. This trial was an organic on-farm trial planted on June 14, 2022. Overhead irrigation was used after planting to promote successful emergence, and the beans were then dry-farmed for the remainder of the season. Plants were pulled from the ground on September 23, 2022, to complete senescence/drydown, and were harvested from the field on September 29, 2022. Samples were placed at -80°C on the day that they were threshed via stationary thresher (one day after harvest). For aim 2, the two dynamic simulated gastrointestinal digestion methods that required less sample mass (∼360 g in total needed for three experimental runs of both the PS and simplified HGS methods vs. an additional ∼360 g needed for three experimental runs of the full HGS method with gastric secretions and emptying) were performed for both varieties from each of Davis (UC Davis Student Farm) and Pescadero, California, using limitedly available seed harvested from organic on-farm trials from the Summer 2019 growing season. The Davis field site was planted on May 30, 2019, furrow-irrigated, cut and windrowed on September 12, 2019, and threshed September 18, 2019. The Pescadero field site was planted on May 24, 2019, used overhead irrigation, uprooted to dry plants on September 14, 2019, and threshed on September 24, 2019. This seed from 2019 was stored at room temperature from harvest until time of analysis.

### 2.3 At-Harvest Trait Analysis

Samples collected in San Gregorio 2022 were harvested from multiple plots within the field. Thus, biological replicates were tested for pre-digestive traits before the seed was pooled to perform digestion tests which required high sample mass (e.g. 120 g of seed per replicate digestion). The samples from the 2019 growing season did not have multiple biological replicates available for testing at-harvest traits, such that only one replicate was examined.

#### 2.3.1 Whole Seed Imaging

Whole beans were imaged (Figure S1) and analyzed to quantify the percent of pigmented seed area in ImageJ as described by Parker et al.*.^49^*In summary, images were taken using a Brother (Nagoya, Japan) DCP-7065DN scanner and then processed using custom macros which were developed in ImageJ (source code available at https://github.com/TravisParker91/Seed-color). The raw data produced from this analysis included the percentage of pigmented area in the scan and the percentage of the image that was seed. The proportion of the seed that was pigmented was calculated by dividing the first trait by the second trait and then multiplied by 100 to represent the percentage of pigmented seed coat area.

#### 2.3.2 At-Harvest Sample Preparation

Whole raw seeds (referred to as raw beans throughout) were ground to a fine powder using a IKA tube mill (TUBE MILL CONTROL S001, Wilmington, NC, USA) with multi-use milling tubes (i.e., grinding chambers; IKA MMT 40.1, Wilmington, NC, USA). Approximately 40 g of sample was split into two subsets (to avoid overloading the grinding chamber), then ground at 25,000 rpm for 78 seconds in 10-second intervals (the final interval was only 8 seconds in duration) and the two ground subsets were then recombined. Ground samples were stored at - 80°C until further analysis.

#### 2.3.3 Phenolic Extractions and Colorimetric Analysis

To extract phenolic compounds from raw seed, 1 g of ground dry bean was added to 5 mL of 50% methanol. Samples were then sonicated in a 2510 Branson water bath (Danbury, CT, USA) for 45 minutes at 24℃, and subsequently placed on a Stovall ‘The Belly Dancer’ shaker (Greensboro, NC, USA) for one hour at room temperature. The extraction was then centrifuged for 5 minutes at 2,103 *g* in a Thermo Scientific Centrifuge (Sorvall Legend XFR, Osterode am Harz, Germany) to pelletize any solid particles. The supernatant was kept for analysis while the pellet was resuspended in a fresh 5 mL aliquot of 50% methanol for further extraction. A second through sixth extraction for each sample was performed by sonicating a fresh suspension for 30 minutes at 24℃, keeping each sequential fraction of supernatant for future analysis. All sample fractions were kept on ice until time of analysis. Extractions were performed in triplicate for each sample using three individual aliquots of ground raw bean. The extractions were then assayed for total phenolic compounds using the Folin–Ciocâlteu reagent as described in section 2.3.3.2, and antioxidant power via Ferric Reducing Antioxidant Power methods as described in section 2.3.3.1 (Figure S2).

##### 2.3.3.1 Antioxidant Power (FRAP)

Antioxidant power was measured in triplicate for each sample via the Ferric Reducing Antioxidant Power (FRAP) method with Trolox used as standard, and was adapted to 96-well microplates to be read on a multi-mode plate reader (Biotek SYNERGY-HTX, Winooski, VT, USA).^53^ Briefly, in each well of the 96-well plate, 190 µL of FRAP reagent plus 10 µL of sample were incubated for 30 minutes under dark conditions until taking the absorbance reading at 593 nm. A standard curve of 50, 100, 200, 300, and 400 mg L^-1^ of Trolox was used to calculate the concentration of Trolox equivalent compounds present in the sample.

##### 2.3.3.2 Total Phenolics (FC)

The total concentration of phenolic compounds was measured in triplicate for each sample via the Folin–Ciocâlteu (FC) method with gallic acid used as standard, and was adapted to 96-well microplates to be read on a multi-mode plate reader.^54^ In each well of the 96-well plate, 80 µL of water, 25 µL of sample, and 5 µL of Folin–Ciocâlteu reagent were incubated for 5 minutes. Then, 80 µL of 7.5% Na_2_CO_3_ was added and the plate was left to incubate for 30 minutes under dark conditions until taking the absorbance reading at 745 nm. A standard curve of 25, 100, 150, 200, and 250 mg L^-1^ of gallic acid was used to calculate the concentration of gallic acid equivalent compounds present in the sample.

#### 2.3.4 At-Harvest Nutritional Quality Traits

Total starch for the ground dry bean samples was determined using a Megazyme Total Starch Assay Kit (AA/AMG, PC: K-TSTA-100A, Wicklow, Ireland) following the AOAC Official Method 996.11,^55^ with minimal adaptation. Specifically, rather than measuring individual reactions in a glass conical tube, the final reaction values were measured at 510 nm by loading 225 µL in triplicate into a 96-well plate to be read on a multi-mode plate reader. This assay was performed in triplicate for each sample using three individual aliquots of ground raw bean. This assay was performed only for the bean samples produced in 2022, as insufficient sample mass was available for the samples produced in 2019.

Ground raw bean samples were also sent to the UC Davis Analytical Lab to determine dry matter, protein, ash, and fat values. Dry matter was measured by oven drying for 3 hours at 105°C. Protein was measured by the AOAC Official Method 990.03,^56^ Protein (Crude) in Animal Feed, via the combustion method. Ash was measured by the AOAC Official Method 942.05,^57^ Ash of Animal Feed. Fat was measured by the AOAC Official Method 2003.05,^58^ Crude Fat in Feeds, Cereal Grains, and Forages. Each trait value was reported as a percentage of seed composition on an “as is” basis (i.e., without having corrected for moisture).

### 2.4 Cooking, Mastication, and Digestion Methods

An overview of the simulated digestion workflow and sampling time points for each digestion method is depicted in Figure 1. All digestive fluids were prepared as described in Sun et al. and formulations provided in Table S2.^6^ All digestive fluids were mixed at room temperature, allowed to stir until all components were fully dissolved, brought to volume, and the pH was adjusted as needed (Table S2) using a Thermo Fisher Scientific AE150 pH meter (Waltham, MA, USA). Water used for digestive fluids was obtained from a Milli-Q Water Purification System (Merck Millipore, Billerica, MA, USA). On the day of a given experiment, all simulated digestion fluids [saliva, gastric juice (GJ), and intestinal fluid (IF)] were warmed to 37°C in a shaking water bath and pH was readjusted as needed (Table S2) prior to digestion.

#### 2.4.1 Cooking Methods

To prepare the cooked beans for each digestion, methods were followed as specified in the variety registration publications for UC Southwest Red and UC Southwest Gold.^51,52^ To prepare the beans, 60 g of dry beans were first soaked in 500 mL of 0.15% NaCl overnight. On the day of the experiment, the soaked beans were rinsed then added to 1.5 L of water for the cooking process. Halfway through the cooking process, 0.9 g of NaCl was added to the beans. For the full HGS method, these volumes were doubled to prepare 120 g of dry beans. The beans were cooked via boiling for 20 minutes followed by simmering for 20 minutes. At the end of this cooking period, the broth was drained and reserved for further analysis. The beans were briefly rinsed in cool tap water to stop the cooking process and used immediately for digestion.

#### 2.4.2 Mastication

Simulated mastication was conducted by first mechanically breaking down samples using the food grinder attachment of a standard KitchenAid mixer (St. Joseph, MI, USA) with a 5 mm plate.^59^ The ground cooked beans were then mixed in a 5 g to 1 mL ratio of food matter (g) to saliva (mL) to reach the final input sample mass needed for each digestion platform, as specified below, for 30 seconds.^59,60^ The bolus was immediately used for the gastric phase of digestion. The remainder of the ground cooked beans were immediately used for analysis of the cooked bean sample.

#### 2.4.3 Gastric Phase

For each digestion, one of the following gastric methods was used for simulation of gastric digestion. These methods were tested as a four-way comparison to determine whether methods with higher throughput and/or requiring smaller sample mass were able to emulate the full HGS method that includes dynamic gastric secretions and gastric emptying.

##### 2.4.3.1 Human Gastric Simulator (full HGS) Method

The HGS was set up and used as previously described to simulate dynamic *in vitro* digestion.^5–8^ The chamber was heated to 37°C, and 35 mL of GJ was added to the HGS bag to simulate fasting gastric fluids.^61^ Next, 200 g of masticated sample was transferred to the HGS bag. The HGS rollers were then turned on at three contractions per minute, and GJ was pumped into the system at a rate of 4.1 mL min^-1^.^62^ Throughout gastric digestion, ∼162 g of sample was taken from the bottom of the stomach bag after 30, 60, 90, and 120 minutes simulating an average gastric emptying rate of 5.4 mL min^-1^. At each time point, the HGS was briefly stopped for sample collection.

For each of the four gastric time points, the solid:liquid ratio of the gastric digesta was determined using a sieve (0.6 mm mesh). The solid:liquid ratio was then utilized to reconstitute two aliquots of approximately 3 g of gastric digesta maintaining the same solid:liquid ratio of the total gastric digesta emptied from the HGS in an opaque 15 mL conical tube for small intestinal digestion.^6^ The remaining solids and liquids were used for analysis of the sample taken at the end of gastric phase (t-120 in Figure 1).

##### 2.4.3.2 Simplified HGS Method

Once the HGS stomach bag was properly fitted to the instrument and the chamber heated to 37°C, 100 mL of GJ was directly added to the bag. Next, 120 g of masticated sample (a 5:1 ratio of cooked sample to saliva) was transferred to the HGS bag, resulting in a 5:6 ratio of GJ to masticated sample. HGS rollers were then turned on at three contractions per minute.^8^ The entire contents of the HGS bag were collected after 120 minutes.^63,64^

As described above (Section 2.4.3.1), the solid:liquid ratio was determined and utilized to reconstitute two aliquots of approximately 10 g of gastric digesta maintaining the same solid:liquid ratio of the total gastric digesta emptied from the HGS in an opaque 50 mL conical tube for small intestinal digestion.^6^ The remaining solids and liquids were used for analysis of the sample taken at the end of gastric phase (t-120 in Figure 1).

##### 2.4.3.3 Peristaltic Simulator (PS) Method

The PS allowed for multiple digestion replicates and/or sample types to be run simultaneously. Each tube was used to digest a single batch of cooked beans. To set up the PS, sample tubes 26 cm in length, made from heat-sealing the bottom of ULINE 4 Mil Poly Tubing (Pleasant Prairie, WI, USA) with an Impulse Sealer (TEW Electric Heating Equipment Co., Type: TISH-300, Taipei, Taiwan), were fitted to the Peristaltic Simulator device for each individual sample type tested. Once the PS chamber was heated to 37°C, 35 mL of GJ was directly added to each fitted tube. Next, 42 g of masticated sample (a 5:1 ratio of cooked sample to saliva) was transferred to each tube resulting in a 5:6 ratio of GJ to masticated sample (same ratio as the simplified HGS method). The device was then turned on to have three peristaltic contractions per minute.^9^ The entire contents of each tube were collected after 120 minutes.^63,64^

As described above, the gastric digesta solids:liquids ratio was determined and two aliquots of gastric digesta were reconstituted as described above and saved for small intestinal digestion. The remaining solids and liquids were used for the analysis of the sample taken at the end of gastric phase (t-120 in Figure 1).

##### 2.4.3.4 Static Digestion Method

For the static gastric digestion method, 35 mL of GJ was directly added to a 250 mL glass bottle. Next, 42 g of masticated sample (a 5:1 ratio of cooked sample to saliva) was transferred to the bottle resulting in a 5:6 ratio of GJ to masticated sample. The bottle was then capped and incubated in a 37°C shaking water bath (TSSWB27, Thermo-Fisher, Waltham, MA, USA) at 100 cycles per minute. The entire contents of the bottle were collected after 120 minutes.^63,64^

As described above (Section 2.4.3.1), the gastric digesta solids:liquids ratio was determined and two aliquots of gastric digesta were reconstituted and saved for small intestinal digestion. The remaining solids and liquids were used for the analysis of the sample taken at the end of gastric phase (t-120 in Figure 1).

#### 2.4.4 Small Intestinal Phase

The gastric digesta samples from all digestion methods underwent the same small intestinal phase. For the small intestinal phase simulation, intestinal fluid (IF) was added in a 4:1 ratio (4 mL IF to 1 g gastric digesta) to each gastric digesta aliquot.^65^ Samples were then incubated in a 37°C shaking water bath (TSSWB27, Thermo-Fisher, Waltham, MA, USA) at 100 cycles per minute. After one hour in the water bath, one of the two samples was removed to represent the 180-minute time point. After two hours, the second sample was removed to represent the 240-minute time point.^66^

### 2.5 Sample Analysis

#### 2.5.1 Moisture content

For the ground cooked beans and for each solid phase gastric digesta sample, three 1.5 g samples were taken for moisture content analysis. Samples were placed in a vacuum oven set to 110°C and dried until constant weight (∼18 hrs). Note, for the full HGS method, moisture content analysis of solid samples was not always possible due to the limited volume of the solids fraction, particularly for the 90- and 120-minute time points. When solid sample volume was limiting, only one or two samples per time point per experimental run were taken for moisture content analysis.

#### 2.5.2 Particle size analysis

For the ground cooked beans and for each solid phase gastric digesta sample, three 1.5 g samples were taken for particle size analysis. The particle size distribution was determined following the procedure by Swackhamer et al.^8^ Images were analyzed using MATLAB R2018b (MathWorks, Natick MA) to obtain the x10, x50, and x90 (the size of a hypothetical sieve through which 10, 50, or 90% of the particle area could pass, respectively) as well as the total number of particles per gram dry weight of sample (Figure S5).^6–8^

#### 2.5.3 pH

For each gastric time point, the pH of the emptied whole digesta was measured using a Thermo Fisher Scientific AE150 pH meter (Waltham, MA, USA).

#### 2.5.4 Sample Preparation for Colorimetric Analyses

##### 2.5.4.1 Broth

The volume of the broth was measured via a graduated cylinder to assess evaporation during cooking. A 10 mL aliquot of broth was then centrifuged for 5 minutes at 2,103 *g* in a Thermo Scientific Centrifuge (Sorvall Legend XFR, Osterode am Harz, Germany). The supernatant was collected and kept on ice until assayed via FRAP and FC methods as described in sections 2.3.3.1 and 2.3.3.2. The remainder of this aliquot was frozen at -20°C for future analysis of starch and protein hydrolysis (as described in section 2.5.5).

##### 2.5.4.2 Cooked Beans

For the cooked sample prior to digestion, 20 g of ground sample was homogenized at 10,000 rpm (Ultra Turrax T18 digital with S18N-19G disperser, IKA Works, Wilmington, NC, USA) for 15 seconds in 20 mL of methanol and left at 4°C for ∼4.5 hours for phenolic compound extraction. Following incubation, the sample was inverted and centrifuged for 5 min at 2,103 *g*. The supernatant was collected and kept on ice until assayed via FRAP and FC methods as described in sections 2.3.3.1 and 2.3.3.2. The remainder of this aliquot was frozen at -20°C for analysis of starch and protein hydrolysis (as described in section 2.5.5).

##### 2.5.4.3 Digesta Sample Preparation

Digesta samples for colorimetric analysis were collected by sampling from the liquid phase of the gastric or intestinal digesta. These samples were immediately mixed in a 1:1 ratio with 7.5% Na_2_CO_3_ to deactivate any gastrointestinal enzymes. Samples were then inverted to mix, centrifuged for 5 minutes at 2,103 *g*, and the supernatant was kept on ice until assayed via FRAP and FC methods as described in sections 2.3.3.1 and 2.3.3.2. Antioxidant power was not measured for small intestinal samples (the 180- and 240-minute time points) due to the coloration of the intestinal fluid interfering with absorbance readings of the plate reader. The remainder of the deactivated sample was frozen at -20°C for analysis of starch and protein hydrolysis (as described in section 2.5.5).

#### 2.5.5 Analysis of Starch and Protein Hydrolysis

Samples stored at -20°C were thawed on ice for approximately 1 hour when ready for analysis of starch and protein hydrolysis.

##### 2.5.5.1 Starch Hydrolysis (Reducing Sugars)

Starch hydrolysis was measured in triplicate for each sample based on the determination of reducing sugars (RDS) following the 3,5-Dinitrosalicylic acid (DNS) assay with maltose used as a standard,^67^ and with adaptation to 96-well microplates.^68^ Under red light conditions, 400 µL of DNS reagent plus 400 µL of sample were mixed, covered with tinfoil, and placed in a boiling water bath for 20 minutes, then on ice for 10 minutes. Samples were then vortexed, and 225 µL of the reaction mixture was added to three wells of a 96-well microplate, and the absorbance was read at 540 nm. If needed, samples were diluted by mixing 100 µL of the reaction mixture with 900 µL of Milli-Q water, and then adding 225 µL of the dilution mixture to three wells of a 96-well microplate to re-measure the absorbance, again at 540 nm. A standard curve of 250, 500, 1000, 1500, and 2000 mg L^-1^ of maltose was used to calculate the concentration of maltose equivalent compounds present in the sample.

##### 2.5.5.2 Protein Hydrolysis (Free Amino Groups)

Protein hydrolysis was measured in triplicate for each sample via the *o*-phthaldialdehyde (OPA) method with glycine used as a standard,^69^ and with adaptation to 96-well microplates.^68^ Briefly, to prepare two solutions with and without OPA (just before measurement), 40 mg of OPA was weighed and added to 1 mL methanol (Methanol-OPA solution). In a volumetric 50 mL flask, 1.25 mL of 20% sodium dodecyl sulfate, 100 μL 2-mercaptoethanol, and 1 mL of Methanol-OPA solution was added and made up to 50 mL with 0.1 M, pH 9.3 sodium tetraborate solution. For the solution without OPA, the Methanol-OPA solution was replaced with pure methanol. After the solutions were prepared, 200 µL of solution containing OPA was pipetted into three wells of a 96-well microplate for each sample. Next, 20 µL of sample was added to each of these wells. After reacting for exactly 4 minutes, the absorbance of the reaction was read at 340 nm. This procedure was then repeated using the solution without OPA for analysis of a sample blank for each sample. A standard curve of 0, 100, 200, 300, and 400 mg L^-1^ of glycine was used to calculate the concentration of NH_2_ groups present in the sample.

### 2.6 Calculations

To calculate the total amount of analyte present in a sample from an individual time point, the value determined by the use of a standard curve was multiplied by the appropriate unit conversion factor and the total liquid volume of the sample; dilution factors were accounted for when applicable due to the addition of intestinal fluid and/or Na_2_CO_3_. These trait values are reported herein as total mg of NH_2_ or total mg of maltose equivalents.

To calculate the amount of nutrient per gram dry weight of bean sample (i.e., mass-standardized values) from an individual time point, the total amount of nutrient present was divided by the dry weight of bean sample that was initially put into the digestion system, with a time point-specific correction for moisture. Specifically, the dry weight of bean sample was calculated by multiplying the known wet weight of cooked bean that was initially put into the system by the percent total solids of the sample, which was determined based on the moisture content of the subsamples taken as described in section 2.5.1. These trait values are reported herein as mg of nutrient per gram dry weight of bean sample.

For the full HGS method, only the total amount of analyte released was calculated due to multiple stages of digestion happening simultaneously (e.g., 90 minutes in the stomach vs. 30 minutes in the stomach followed by 60 minutes in the small intestine; Figure S8). Starch and protein hydrolysis were determined via the calculations described in Sun et al. for both individual time points and cumulative absorption throughout gastric and small intestinal digestion.^6^ In summary, for each sampling time point, the total amount of nutrient present in the liquid sample was calculated as described above. Two steps were then needed to calculate the absorption profile of the nutrient (i.e., the total amount of nutrient released at each 30-minute increment of the digestive process). First the differences between intestinal time points generated from the same gastric digesta sample were calculated, including the difference between the gastric digesta sample itself and the first 30-minute intestinal sample. Then, using these calculated differences for the intestinal time points in conjunction with the previously obtained gastric values, samples with equivalent total digestion time (i.e., the cumulative time spent in both the gastric and intestinal phases) were summed to obtain the value of total analyte released at each 30-minute increment of the digestive process. A cumulative release curve was then generated by summing each individual time point of the absorption profile. Final values were reported as total mg of NH_2_ or total mg maltose equivalents released at the given time point (Figure S4). The values at the 120-, 180-, and 240-minute time points (based on total digestion time) from these results were used for the four-way digestion method comparison, as these were the same time points sampled in the other three digestion methods. Cumulative release curves (and the resulting trait values) for total phenolics and antioxidant power were not calculated for the full HGS due to the potentially non-cumulative nature of these compounds.

### 2.7 Statistical Analyses

For the comparison of the four digestion platforms tested in aim 1, a mixed analysis of variance (ANOVA) was conducted using RStudio (RStudio, 2021.09.1, Build 372, © 2009-2021 RStudio, PBC). Specifically, a model was fitted using the *lm()* function from the lme4 package and included the main and interaction effects of digestion method and time point for all traits except pre-digestion total phenolics and antioxidant power, for which the statistical analyses are described in the next paragraph.^70^ Group means for each digestion method-time point combination were then calculated using the *emmeans()* function from the emmeans package version 1.7.0.^71^ A post-hoc Tukey’s HSD (Honestly Significant Difference) test was then conducted using the *cld*() function from the emmeans package to compare means at a significance level of ɑ = 0.05 (Supplemental File S1). Likewise, for the comparison of the beans tested in aim 2, ANOVA was conducted by fitting a model including the main and interaction effects of genotype, growing location, digestion method, and time point using the *lm()* function for all traits except pre-digestion total phenolics and antioxidant power, for which the statistical analyses are described in the next paragraph. Group means were then calculated and followed by a post-hoc test as described above (Supplemental File S1). In both analyses, a post hoc test was only conducted for the higher-order term in which a given variable was involved; e.g., if an interaction term was significant, a post hoc test was conducted for only the interaction (and not for the main effects) so as not to interpret main effects in the presence of an interaction.

For pre-digestion total phenolics and antioxidant power, mixed ANOVA and post hoc tests were only applied within time points (i.e., raw beans, cooked beans, or broth) as extraction methods differed for each time point. Additionally, post hoc tests were only applied to the genotype by environment interaction term and not to main effects due to data structure imbalance (i.e., so that all group means were estimable even with Southwest Red not having been tested in San Gregorio) (Supplemental File S1). The UC Southwest Gold sample produced in San Gregorio (used in Aim 1) was only compared to the UC Southwest Gold and Red samples produced in Pescadero and Davis in 2019 (used in Aim 2) for pre-digestion total phenolics and antioxidant values (and percentage of pigmented seed coat area); this comparison was not conducted for any other trait or time point.

Averages and standard deviations were calculated across replicates when applicable. For the statistical comparison on pre-digestive traits, a two-tailed unpaired t-test was applied.

## 3 Results

### 3.1 At-Harvest Traits (raw beans)

#### 3.1.1 Seed coat coloration/patterning

For UC Southwest Gold samples grown in 2019, 63.7% (± 1.8%, SD) and 9.4% (± 0.9%, SD) pigmented seed coat area were observed for Davis and Pescadero, respectively (Table S3). For UC Southwest Red samples grown in 2019, 70.8% (± 0.8%, SD) and 21.0% (± 5.7%, SD) pigmented seed coat area were observed for Davis and Pescadero, respectively (Figure 2). UC Southwest Gold grown in San Gregorio in 2022 displayed a phenotype similar to UC Southwest Gold grown in Pescadero in 2019 with 17.7% (± 1.0%, SD) pigmented seed coat area (Figure S1). This trait was found to be significantly different (*P*-value = 0.002) between the Davis and coastal (i.e. San Gregorio and Pescadero) locations.

**Figure 2.**
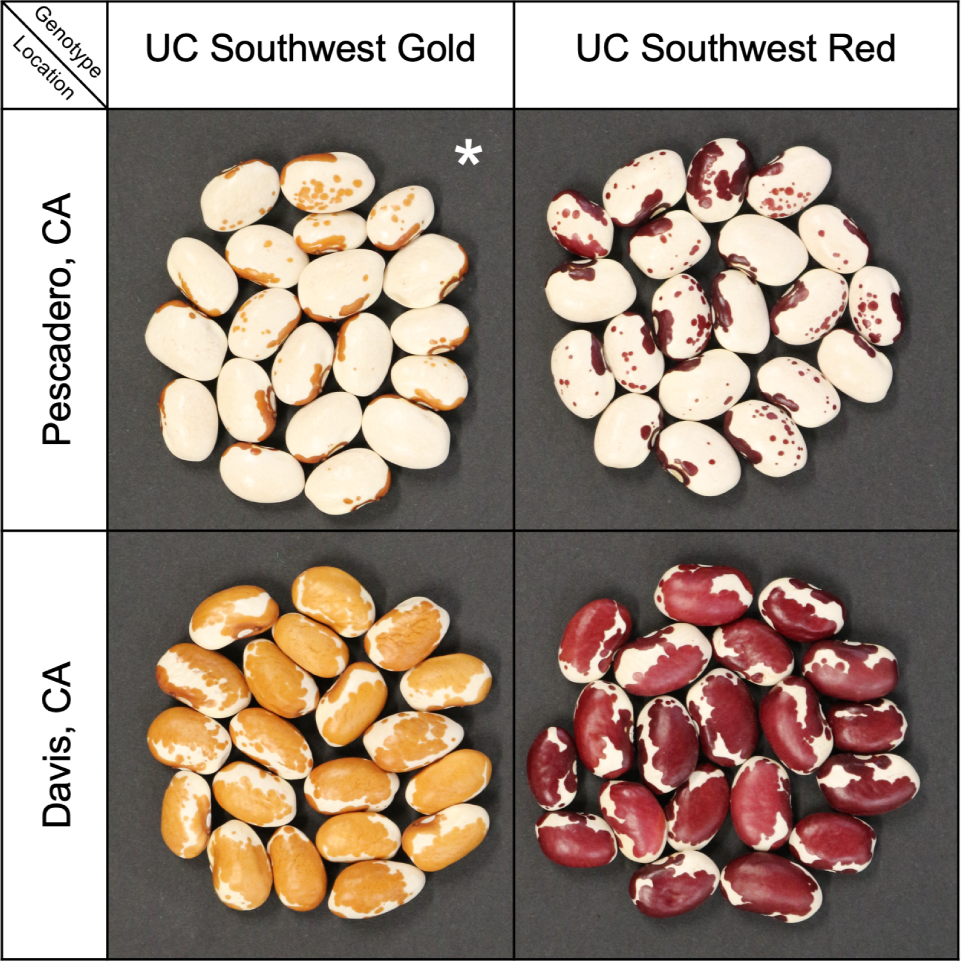
Seed of the UC Southwest varieties, depicted based on the growing locations in 2019 in which these samples were produced. Adapted from Parker et al..^49^ *UC Southwest Gold grown in San Gregorio in 2022 displayed a phenotype similar to UC Southwest Gold grown in Pescadero in 2019 (Figure S1).

#### 3.1.2 Nutritional Quality Traits

Values for nutritional quality traits measured on raw beans are reported in Table 1. The bean samples from coastal growing locations had significantly higher (*P*-value = 0.038) protein levels than beans produced in Davis. Post hoc tests for comparison of group means were otherwise not feasible for these traits at this time point due to only one biological replicate being available for the 2019 samples. For the coastal growing locations, UC Southwest Gold bean samples grown in San Gregorio in 2022 had comparable values for crude fat, slightly lower values for protein, and slightly higher values for moisture and ash compared to both the UC Southwest Gold and UC Southwest Red bean samples grown in Pescadero in 2019; however, no significant differences were detected for these traits across genotypes and growing environments. For the beans grown in 2019 in Davis, the UC Southwest Gold bean samples had slightly lower values for protein content compared to the UC Southwest Red bean samples; likewise, these differences were not statistically significant.

**Table 1.** Nutritional quality trait values in ground raw bean samples. Values are reported for moisture, crude fat, starch, protein, total nitrogen, and ash (total mineral) percentages. Each value is reported as a percentage of seed composition on an “as is” basis (i.e., without having corrected for moisture). Averages and standard deviations across biological replicate samples are reported when applicable (n = 3 for UC Southwest Gold, San Gregorio, 2022). NAs are reported for Starch (%) in the 2019 samples as sample mass was limiting.

| Genotype | Location | Year | Moisture (%) | Crude Fat (%) | Starch (%) | Protein (%) | Ash (%) |
| --- | --- | --- | --- | --- | --- | --- | --- |
| UC Southwest Gold | San Gregorio, CA | 2022 | 10.2 ± 0.1 | 1.2 ± 0.1 | 28.3 ± 0.5 | 16.8 ± 0.2 | 4.39 ± 0.14 |
| UC Southwest Gold | Pescadero, CA | 2019 | 9.0 | 1.2 | NA | 18.3 | 4.26 |
| UC Southwest Gold | Davis, CA | 2019 | 9.1 | 1.1 | NA | 20.2 | 4.17 |
| UC Southwest Red | Davis, CA | 2019 | 8.9 | 0.9 | NA | 21.8 | 4.13 |
| UC Southwest Red | Pescadero, CA | 2019 | 9.1 | 1.2 | NA | 18.4 | 4.12 |

### 3.2 Pre-Digestive Phenolics and Antioxidant Power

The effects of genotype, growing location, and their interaction were significant (all *P*-values < 0.001, except the interaction term had a *P*-value of 0.004 for total phenolics) for both total phenolics and antioxidant power in the raw bean samples (Supplemental File S1). For total phenolics in the raw bean samples, UC Southwest Red grown in 2019 in Davis showed significantly higher values compared to all other genotype-environment combinations (Supplemental File S1 and Figure 3, panel A). Additionally, UC Southwest Gold grown in 2019 in Davis showed significantly higher values compared to the beans grown in coastal locations (Pescadero and San Gregorio). Over the six rounds of extraction used to separate total phenolics from the raw bean samples, 77.8 to 87.0% of the total phenolics extracted came out within the first three rounds of extraction (Figure S2). The sixth round of extraction produced approximately 3.1 to 6.8% of the total phenolics extracted. When calculating a ratio of the percentage of pigmented seed coat area and total phenolics (mg gallic acid equivalents per gram dry weight sample) in raw beans, this ratio had a value of 36.8 and 39.8 for UC Southwest Red and Gold in Davis, 15.1 for UC Southwest Red in Pescadero, and 7.1 and 13.6 for UC Southwest Gold in Pescadero and San Gregorio (respectively). For antioxidant power in the raw bean samples: again, UC Southwest Red grown in 2019 in Davis showed significantly higher values compared to all other genotype-environment combinations (Supplemental File S1 and Figure 3, panel B). Additionally, both genotypes grown in 2019 in Pescadero showed significantly lower values compared to all other genotype-environment combinations.

**Figure 3.**
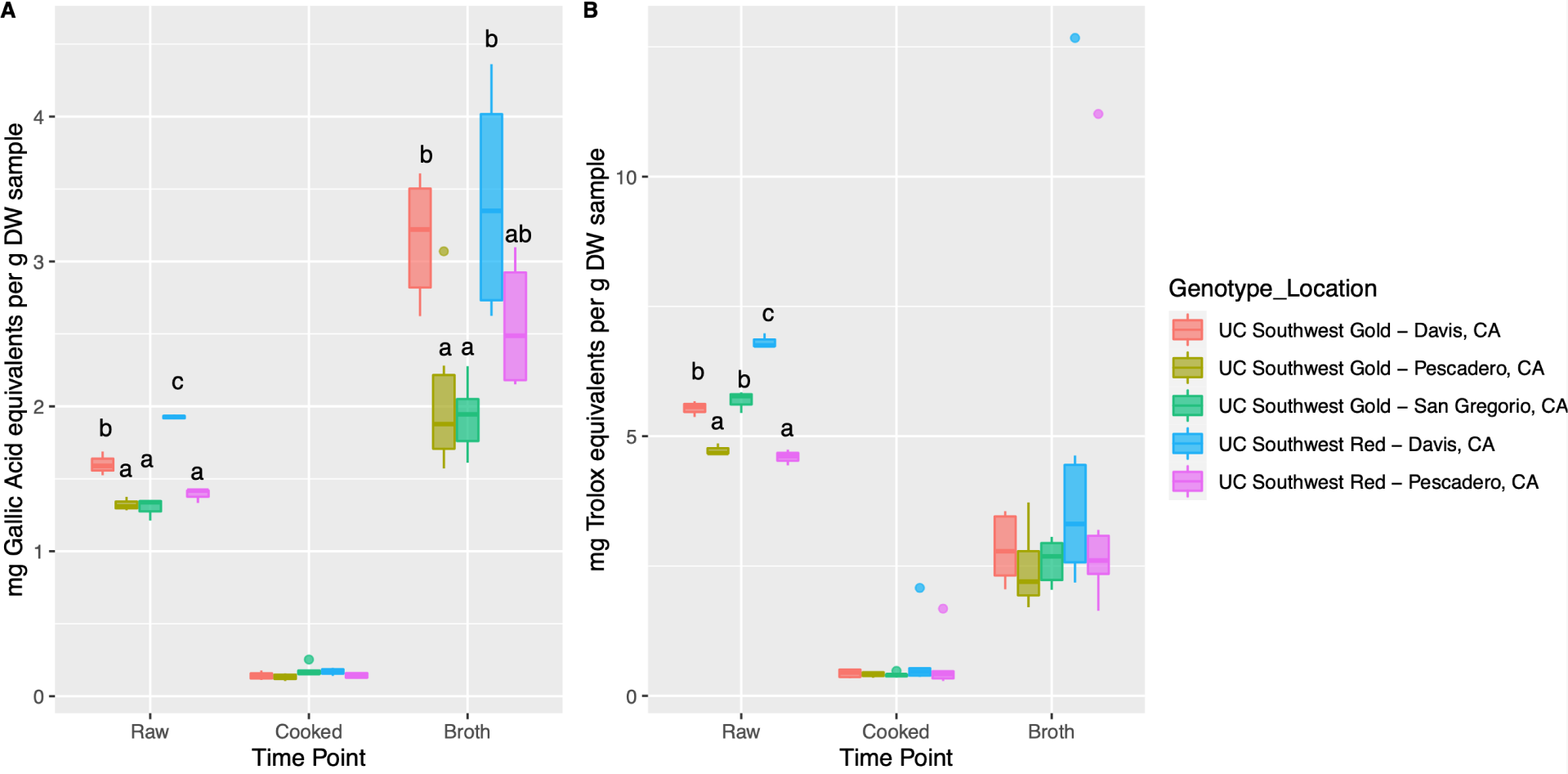
Levels of total phenolics and antioxidant power in the raw bean samples, cooked bean, and broth. A) Total phenolics were determined via FC assays. B) Antioxidant power was determined via FRAP assays. Significance was determined using a Tukey’s Honestly Significant Difference post hoc test following mixed ANOVA. Comparisons were made within rather than across time points, as differing extraction methods were used for each time point displayed. Letters were not displayed for time points within which no significant differences were observed.

The effect of growing location (*P*-value = 0.0095) was the only significant factor for total phenolics in the pre-digestion cooked bean samples (Supplemental File S1). For antioxidant power in the cooked bean samples, no terms were found to be significant. The effects of genotype (*P*-value = 0.005) and growing location (*P*-value < 0.001) were significant for total phenolics in the broth samples (Supplemental File S1). Both genotypes grown in 2019 in Davis had significantly higher values for total phenolics in the broth samples compared to UC Southwest Gold grown in coastal locations (Supplemental File S1 and Figure 3, panel A). For antioxidant power in the broth samples, no terms were found to be significant.

### 3.3 Four-way Digestion Comparison

Of the four digestion methods tested, samples from the full HGS and the simplified HGS method had significantly higher trait values than the PS and static digestion methods for at least one digestive time point for three of the four measured traits (antioxidant power and starch and protein hydrolysis; Figures 4 and 5). Overall, the general trend was that the highest trait values were observed from the full HGS method, followed by the simplified HGS method, PS method, and lastly, the static digestion method (which had the lowest observed values).

**Figure 4.**
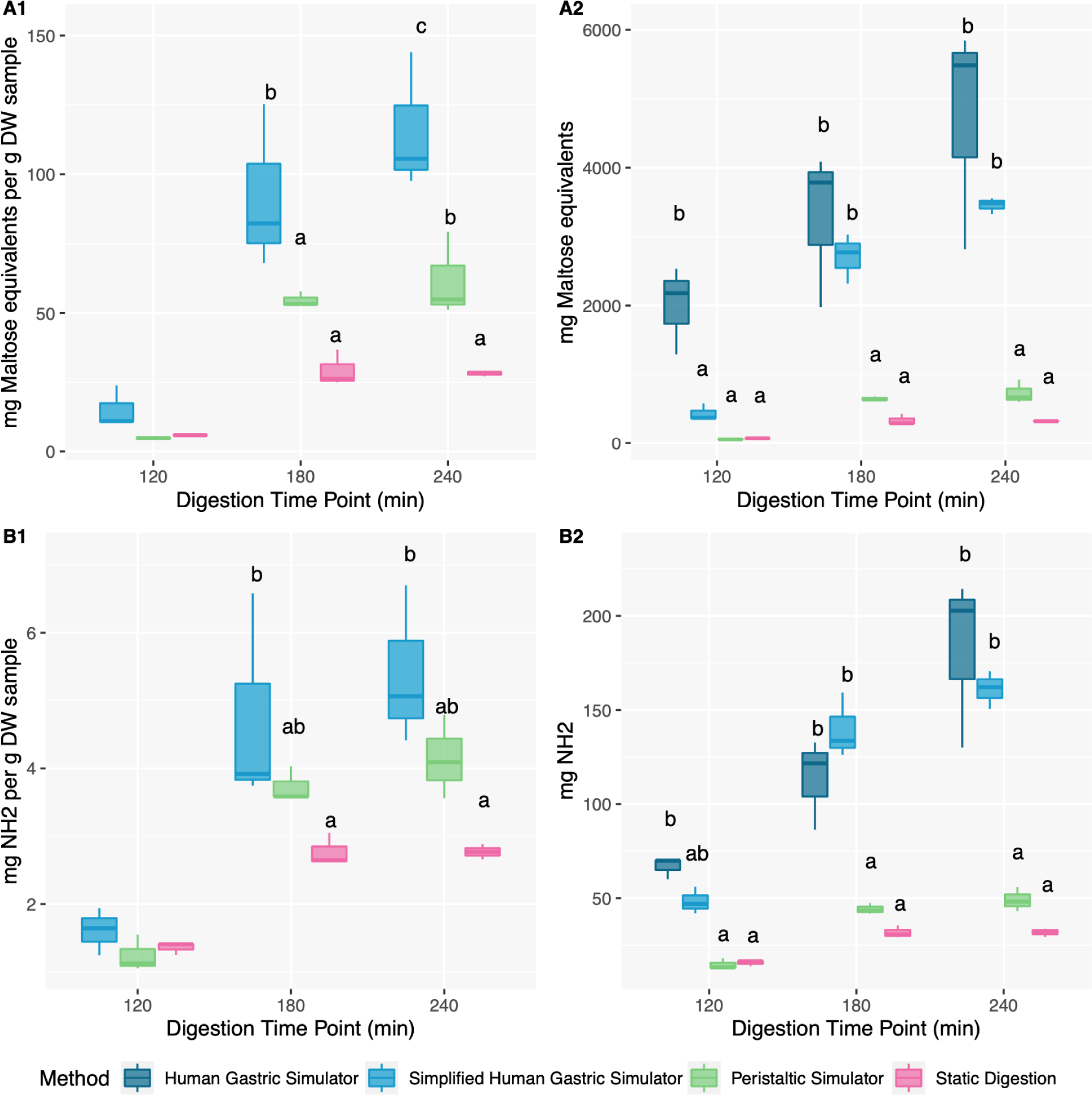
Starch and protein hydrolysis during the digestive time points for the four-way simulated digestion method comparison. The 120-minute time point represents the end of the gastric phase of digestion, and the 180- and 240-minute time points represent one and two hours of small intestinal digestion, respectively. A) Starch hydrolysis was determined via RDS assays. B) Protein hydrolysis was determined via OPA assays. Significance was determined using a Tukey’s Honestly Significant Difference post hoc test following mixed ANOVA. Comparisons were made within time points and letters were not displayed for time points within which no significant differences were observed.

**Figure 5.**
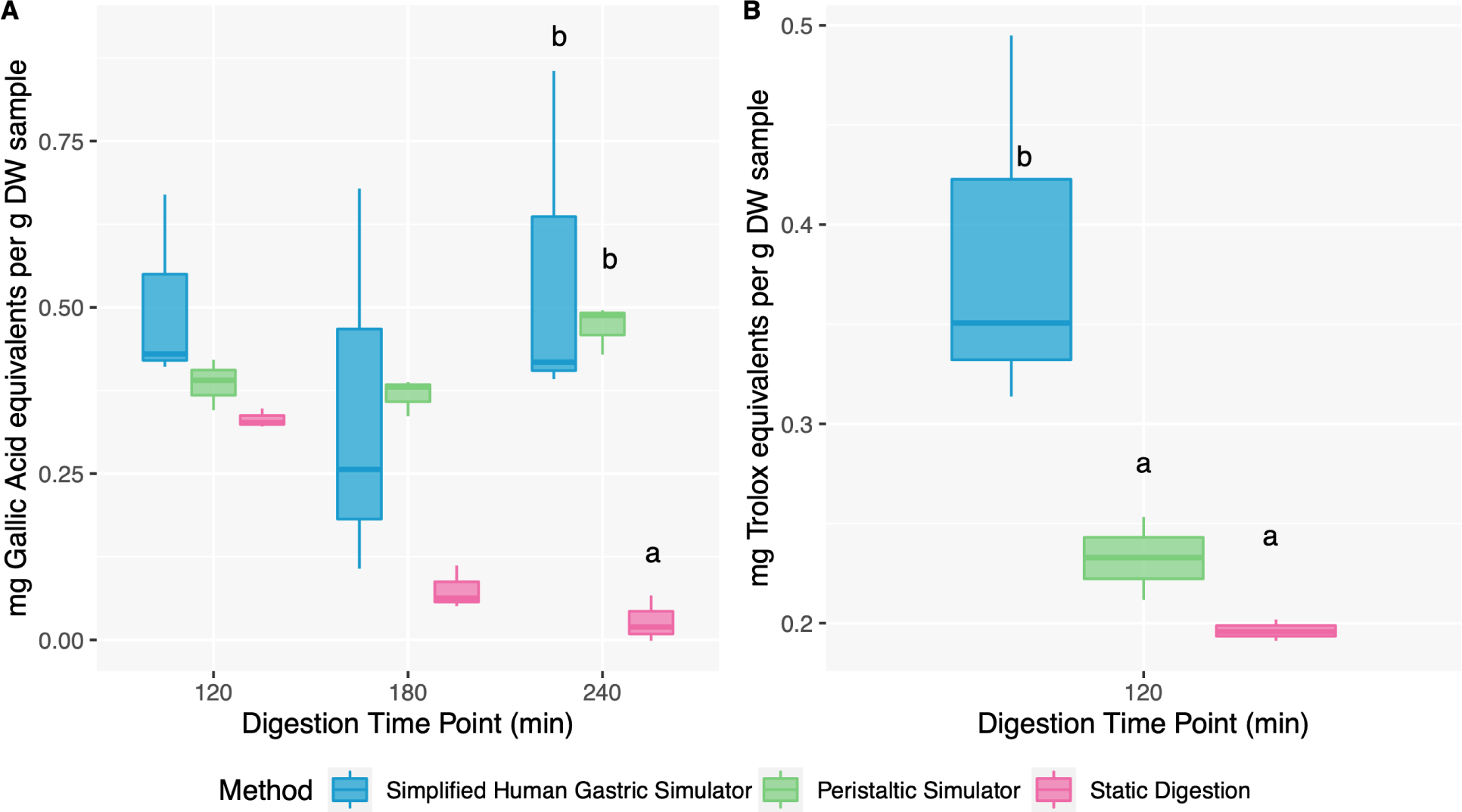
Levels of total phenolics and antioxidant power during the digestive time points for the simulated digestion method comparison. The 120-minute time point represents the end of the gastric phase of digestion, and the 180- and 240-minute time points represent one and two hours of small intestinal digestion, respectively. A) Total phenolics were determined via FC assays. B) Antioxidant power was determined via FRAP assays. Significance was determined using a Tukey’s Honestly Significant Difference post hoc test following mixed ANOVA. Comparisons were made within time points and letters were not displayed for time points within which no significant differences were observed.

The effects of digestion method, time point, and their interaction were all significant for both mass-standardized and non-mass standardized starch hydrolysis during digestion (all *P*-values < 0.001, except the interaction term had *P*-values of 0.003 for mg of maltose equivalents per gram dry weight bean sample and 0.005 for total mg of maltose equivalents) (Supplemental File S1). The effects of both digestion method and time point were significant (all *P*-values < 0.001) for both mass-standardized and non-mass standardized protein hydrolysis, and the interaction of method and time point was also significant (*P*-value < 0.001) for non-mass standardized protein hydrolysis (Supplemental File S1). For all dynamic digestion methods, starch and protein hydrolysis increased over the course of digestion (Figure 4). Mass-standardized starch hydrolysis (i.e., mg of maltose equivalents per gram dry weight bean sample) (Figure 4, panel A1) was significantly higher for the simplified HGS method compared to the PS and static methods for the last two time points and was also significantly higher for the PS method compared to the static method for the final time point. Mass-standardized protein hydrolysis (i.e., mg of NH_2_ per gram dry weight bean sample) (Figure 4, panel B1) was significantly higher for the simplified HGS method compared to only the static method for the last two time points. However, the non-mass standardized results for starch hydrolysis (total mg of maltose equivalents) and protein hydrolysis (total mg of NH_2_) (Figure 4, panels A2 and B2) indicated that while the full and simplified HGS methods had significantly higher values compared to the PS and static methods for the last two time points, they were not significantly different from each other. For total mg of maltose equivalents, the full HGS method was significantly higher than the other three digestion methods at the 120-minute gastric time point. For total mg of NH_2_, the full HGS method was significantly higher than the PS and static methods at the 120-minute gastric time point, but the simplified HGS method was not significantly different from any of the other three methods.

Reducing sugars and free amino groups were also observed in the broth. While the levels of free amino groups in the broth were fairly comparable to those during the course of digestion, the levels of reducing sugars in the broth were much lower than those during the digestive process. The effect of time point (cooked bean vs. broth) was significant for reducing sugars and free amino groups per gram dry weight of sample (all *P*-values < 0.001); namely, both of these traits were significantly lower in the cooked bean samples (i.e., at the zero-minute time point) than in the broth (Figure S7, Supplemental File S1). Total reducing sugars and free amino groups (i.e., total mg of maltose equivalents and NH_2_) were higher in the broth for samples being inputted to the full HGS (for which a larger sample mass was used) than for samples being inputted to the other three gastric digestion methods (all *P*-values < 0.001).

The effect of digestion method was significant (*P*-value < 0.001) for total phenolics during digestion (Supplemental File S1). Total phenolics were significantly higher with the simplified HGS and PS methods than the static method for the last (240-minute) time point assayed (Figure 5). The effect of digestion method (*P*-value = 0.014) was significant for antioxidant power, which was significantly higher for the simplified HGS at the 120-minute gastric time point compared to the PS and static methods (Figure 5, Supplemental File S1). Due to the complex nature of phenolic and other antioxidant formation and breakdown (such that their levels are not cumulative over time), these traits were not calculated for the full HGS as the calculations for that digestion method require stability of the measured compound (or analyte) once formed.

For the full HGS method, the pH of the emptied gastric digesta started at 3.27 (± 0.778, SD) after 30 min, rose to 3.44 (± 0.248, SD) after 60 min, before lowering to 2.71 (± 0.699, SD) and ending at 1.60 (± 0.092, SD) for the 90- and 120-minute time points, respectively (Figure S3). The cumulative values (across the eight time points at 30-minute increments; Figure S4) for total hydrolyzed starch and protein were calculated to be 14,611 mg of maltose equivalents (± 6,141 mg, SD) and 678 mg of NH_2_ (± 218 mg, SD) respectively. The amount of starch and protein hydrolysis measured at each individual time point rose as gastric digestion progressed, peaking at 120 minutes for both traits (4089 mg of maltose equivalents (± 1100 mg, SD) and 231 mg of NH_2_ (± 39.5 mg, SD), respectively) (Figure S4). However, over the course of the small intestinal digestion phase, the amount of starch and protein hydrolysis measured at each individual time point became progressively less. Particle size analysis of gastric digesta indicated a higher number of particles present in gastric samples collected during the full HGS method (when collection was possible at 30, 60, and 90 minutes), indicating a higher degree of particle breakdown for this digestion method (Figure S5). The 120-minute time point sample from the full HGS method was limited and prioritized for other analyses, therefore statistical comparison of particle size for the gastric time point between this and the other digestion methods was not feasible (due to the other digestion methods only being sampled at 120 minutes for the gastric phase).

### 3.4 UC Southwest Red/Gold Digestion Comparison

The effects of digestion method and time point were significant (all *P*-values < 0.001) for both mass-standardized and non-mass standardized results for mg of maltose equivalents and NH_2_ during digestion in this comparison between bean samples (Supplemental File S1). The simplified HGS exhibited significantly higher values than the PS for all four of these starch and protein traits. As was also observed in the four-way digestion method comparison, starch and protein hydrolysis increased over the course of digestion for both digestion methods tested (Figure 6). For both total and mass-standardized results for mg of maltose equivalents and for total mg of NH_2_, trait values were significantly different at each of the three digestive time points, with the 120-minute time point showing significantly lower values than the 180-minute time point, and both of these time points showing significantly lower values than the 240-minute time point (Supplemental File S1). For mass-standardized protein hydrolysis (i.e., mg of NH_2_ per gram dry weight beans), the 120-minute time point was significantly lower than the two small intestinal (180- and 240-minute) time points, which were not significantly different from each other. Additionally, the main effects of genotype (*P*-value = 0.026) and environment (*P*-value = 0.030) were significant for total mg of NH_2_ during digestion, and the interaction of genotype and environment was significant (*P*-value = 0.021) for total mg of maltose equivalents during digestion. For total mg of NH_2_, Davis showed significantly higher values than Pescadero, and UC Southwest Red showed significantly higher values than UC Southwest Gold (Supplemental File S1). For total mg of maltose equivalents, no genotype-environment combinations exhibited significant differences.

**Figure 6.**
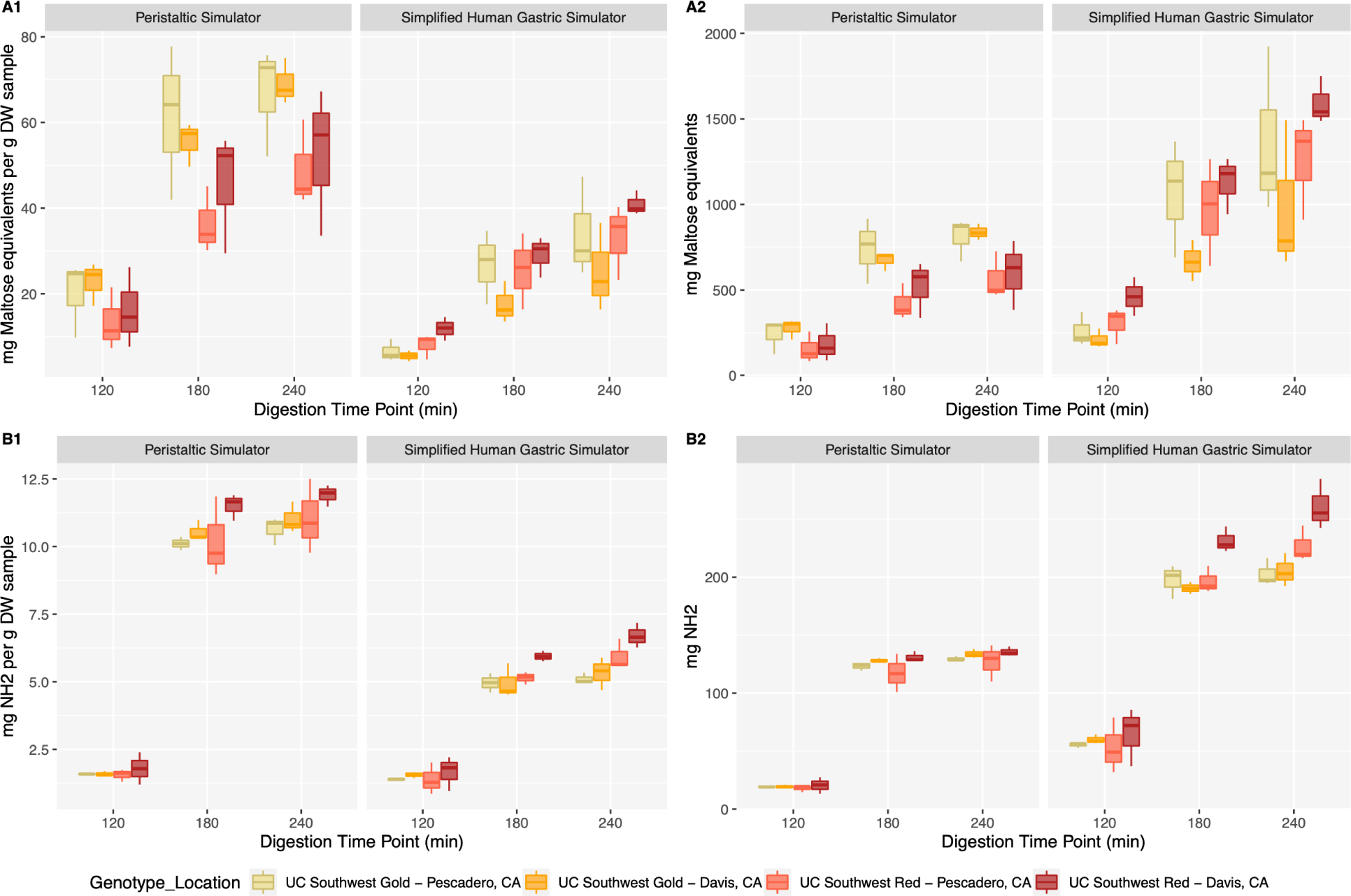
Starch and protein hydrolysis during the digestive time points for the UC Southwest Red and Gold variety comparison using both the simplified HGS and PS digestion methods. The 120-minute time point represents the end of the gastric phase of digestion, and the 180- and 240-minute time points represent one and two hours of small intestinal digestion, respectively. A) Starch hydrolysis was determined via RDS assays. B) Protein hydrolysis was determined via OPA assays. Significance was determined using a Tukey’s Honestly Significant Difference post hoc test following mixed ANOVA. No significant differences were found between genotype-environment combinations in post hoc tests, such that connecting letters are not depicted in this plot.

Reducing sugars and free amino groups were also observed in the broth from these samples, though levels observed from the cooked bean samples were low (Figure S6). The effects of both digestion method and time point (cooked beans vs. broth) were significant (all *P*-values < 0.001) for reducing sugars (observed in both digestion methods for both total mg of maltose equivalents and mg of maltose equivalents per gram dry weight of bean sample) in the cooked bean and broth samples. For free amino groups, however, the effects of genotype (*P*-values = 0.011 and 0.006, respectively), growing location (*P*-values = 0.002 and 0.005, respectively), digestion method (*P*-values = 0.009 and < 0.001, respectively), and time point (both *P*-values < 0.001) were all significant for both total mg of NH_2_ and mg of NH_2_ per gram dry weight of bean sample. The broth from UC Southwest Red samples produced in Davis in 2019 showed significantly higher levels of free NH_2_ (observed in both digestion methods for both total mg of NH_2_ and mg of NH_2_ per gram dry weight of bean sample) than broth from UC Southwest Gold samples produced in Pescadero in 2019 (Figure S6). Significant differences were not observed in the broth for any other pairwise comparison of samples for free amino groups, nor for any pairwise comparisons for reducing sugars.

For total phenolic compounds released during the course of digestion (Figure 7), the effects of time point (*P*-value < 0.001) and environment (*P*-value = 0.035) were significant for total phenolics (Supplemental File S1). Significantly higher phenolics were observed from samples produced in Davis than those produced in Pescadero. Total phenolics were significantly higher at the 180-minute than the 120- or 240-minute time points, which were not significantly different from each other. However, all values were extremely low (< 0.5 mg gallic acid equivalents per gram dry weight beans) in comparison to the levels of total phenolic compounds released into the broth (ranging from ∼1.5 to 4.5 mg gallic acid equivalents per gram dry weight beans) (Figures 3 and 7). Likewise, antioxidant power was lower in gastric samples (< 0.75 mg Trolox equivalents per gram dry weight beans) in comparison to the broth samples (> 2 mg Trolox equivalents per gram dry weight beans) (Figures 3 and 7). The effect of digestion method was significant (*P*-value < 0.001) for antioxidant power during the 120-minute gastric time point, with significantly higher values observed in the PS than the simplified HGS (Supplemental File S1).

**Figure 7.**
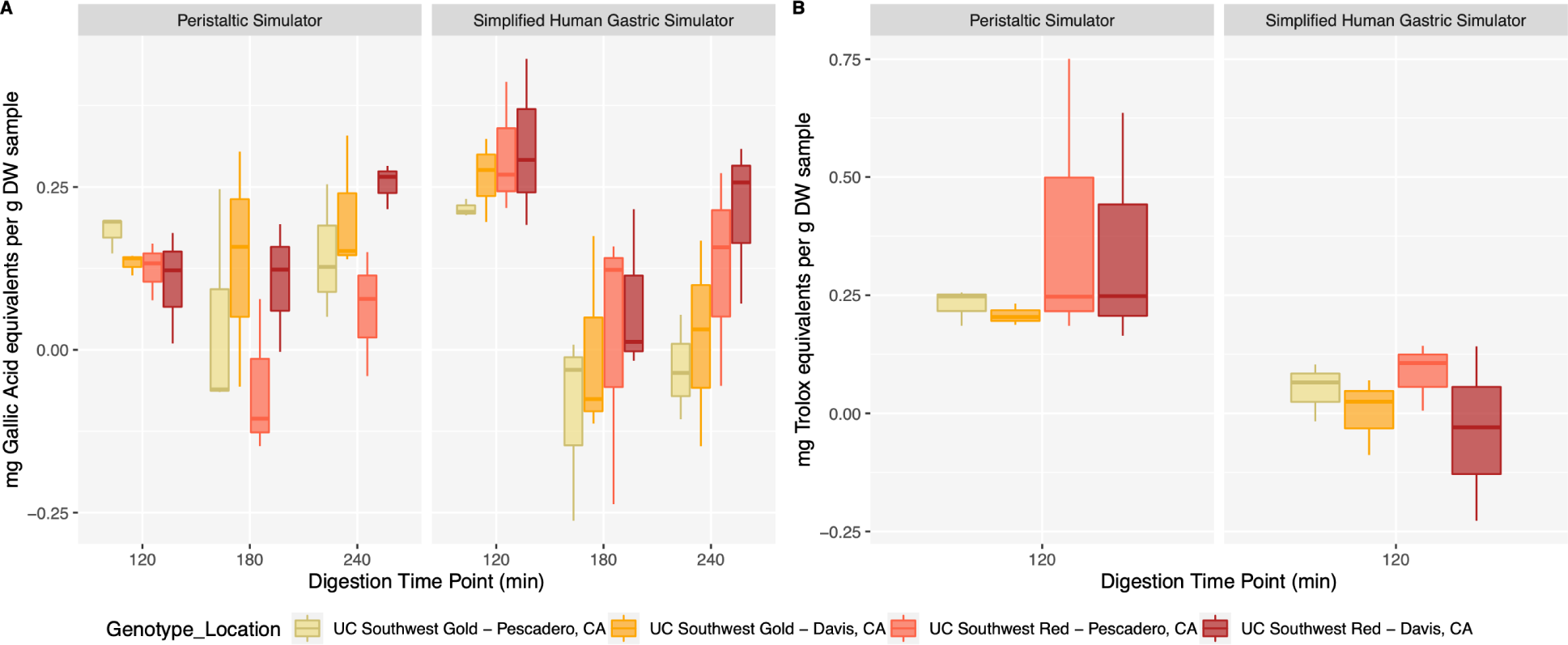
Levels of total phenolics and antioxidant power during the digestive time points for the UC Southwest Red and Gold variety comparison using both the simplified HGS and PS digestion methods. The 120-minute time point represents the end of the gastric phase of digestion, and the 180- and 240-minute time points represent one and two hours of small intestinal digestion, respectively. A) Total phenolics were determined via FC assays. B) Antioxidant power was determined via FRAP assays. Significance was determined using a Tukey’s Honestly Significant Difference post hoc test following mixed ANOVA. No significant differences were found between genotype-environment combinations in post hoc tests, such that connecting letters are not depicted in this plot.

## 4 Discussion

### 4.1 Pigmented Seed Coat Area and Pre-digestion Total Phenolics

Significant differences in the percentage of pigmented seed coat area by environment were observed from image analysis (Table S3), as could be expected based on the visual appearance of the seeds (Figure 2) and as also found in Parker et al. for these genotypes in a partially overlapping set of environments.^49^ The beans grown in Davis had significantly more pigmentation and total phenolics in raw beans compared to those grown in Pescadero. Previous studies have also found higher total phenolics in beans with colored seed coats than those with white seed coats.^32,72^ The beans grown in Davis also had a higher ratio of pigmented seed coat area to total phenolics than those in Pescadero (36.8 and 39.8 vs 15.1 and 7.1, respectively)—i.e., percentage of pigmented seed coat area did not correspond 1:1 with total phenolics concentrations, which may inform breeding efforts for one or both of these traits.

High values for total phenolics were observed in the broth, which is consistent with previous studies.^73–75^ Phenolic extraction for both the raw and cooked bean samples could be improved upon to maximize comparability of phenolic values within pre-digestive time points and between pre-digestive and digestive time points. Although 77.8 to 87.0% of the phenolics were extracted within the first three rounds of extraction (and previous studies have utilized three rounds of extraction ^76^), non-negligible levels of phenolics (3.1 to 6.8%) were still being extracted from the raw beans even in a sixth round of extraction (Figure S2). Based on these results, it would be advisable to test multiple rounds of extraction for quantification of total phenolics even when using published protocols. It is likely that the values seen from the cooked beans were particularly low due to the standard extraction method used during the ongoing digestions (which were conducted prior to optimization of the protocol for raw beans). Additionally, the FC and FRAP assays were conducted during digestion (i.e., on the day of a given experimental run), such that only one round of extraction for use in the FC and FRAP assays was feasible due to the need to execute steps in the digestion workflow in the requisite timing. While significant differences between genotype-environment combinations were observed for antioxidant power in raw beans (with means ranging from 4.6 to 6.8 mg Trolox equivalents per gram dry weight sample; Figure 3, Supplemental File S1), significant differences were not observed in cooked beans nor broth. This relationship between pre-digestive traits would merit further examination in bean samples with a broader range of antioxidant power, e.g. in a larger and more diverse set of cultivars as observed in Madrera et al.^72^

### 4.2 Nutrient Bioaccessibility in Contrasting Sample Types

Even with a small number of unique sample types in aim 2, differences were observed between genotypes for one trait (protein hydrolysis) and between environments for two traits (total phenolics and protein hydrolysis) at specific digestive time points (Supplemental File S1). In this study, samples with higher levels of protein hydrolysis during digestion also had higher protein levels in raw beans (Table 1). Characterizing the impacts of protein-phenolic and starch-phenolic physical/biochemical interactions on protein and starch hydrolysis during digestion are of potential interest for future studies, as these interactions could impact hydrolysis even in samples that have the same protein levels in raw beans.^38–41^ The beans grown in Davis maintained an overall higher level of total phenolics during digestion (Supplemental File S1). Cárdenas-Castro et al. found that two common bean varieties had significantly different levels of total phenolics in the cooked mashed stage, but significant differences between those varieties were not observed in the levels extracted in the gastric and small intestinal phases (rather only in the soluble and insoluble indigestible fractions).^77^ No significant differences between genotype-environment combinations were observed for antioxidant power at the 120-minute time point (when values were near zero), which again would be helpful to examine in cooked bean samples representing a broader range of antioxidant power.

Interestingly, trends in total phenolics over the course of digestion were not consistent (e.g., phenolics were observed to be lowest at the 180-minute time point most commonly but trends of increasing or decreasing over the course of digestion were also observed), which could be illustrative of the potentially non-cumulative nature of phenolics through digestion.^78^ For example, phenolics were substantially lost from chokeberry juice during *in vitro* intestinal digestion,^79^ which was consistent with findings in other plant-based foods.^80^ Bermúdez-Soto et al. and Li et al. determined that such losses were due primarily to the mildly alkaline conditions in the small intestine rather than interactions with digestive enzymes based on the *in vitro* digestion and recovery of pure standards (which would not have accounted for food matrix effects particularly for a solid food),^79,81^ with potential for phenolic derivatives to form.^82^ Sancho et al. found total phenolics extracted from raw common bean decreased after (compared to before) *in vitro* gastointestinal digestion in a shaker.^83^ While a minor but insignificant decrease was observed in the static digestion method herein (0.332 to 0.075 to 0.028 mg gallic acid equivalents per gram dry weight bean sample at 120, 180, and 240 minutes; Figure 5), such a trend was not observed for the PS and simplified HGS. Further examination of the abundance, *in vivo* activity, and turnover of individual phenolic compounds throughout dynamic digestion would be informative in maximizing the benefits (and minimizing the detrimental effects) of bean varieties for human nutrition.^35^

### 4.3 Diversity in Dynamic Digestion Models

The results of this study are consistent with earlier studies, which showed that more complex dynamic digestion models are indeed better for simulating human digestion.^5^ However, it was found herein that even simpler dynamic digestion models with higher throughput or a reduced sample mass requirement gave rise to more nutrient release than a static simulated digestion method. The simplified HGS method notably was only significantly different from the full HGS for non-mass standardized starch hydrolysis at the 120-minute gastric time point (out of the three digestive time points assayed for non-mass standardized protein and starch hydrolysis). Both the simplified HGS and PS methods had overall lower standard error in comparison to the full HGS method, namely due to the complexity of extracting samples at multiple intermediate gastric time points from the full HGS both for analysis and for real-time transfer to the small intestinal phase. This simplified HGS method could be of value to future simulated digestion experiments given its reduced sample mass requirement and reduced protocol complexity. Previous studies have compared variations of the full HGS method ^84^ and comparisons of other dynamic models that include gastric emptying to static models.^85^ However, models that include gastric emptying are not suited for limited sample masses. Additionally, Barros et al. compared a dynamic model that did not include gastric emptying to a static model.^86^ While there is vast diversity in digestion method protocols, few direct comparisons have been made between the presence or absence of gastric emptying within the same dynamic model (e.g., the full vs. simplified HGS methods) and between dynamic models that can accommodate different sample masses (e.g., the full HGS, simplified HGS, and PS methods).^87,88^

The PS shows promise as a dynamic digestion platform whose throughput is approximately the same as that of the static model considering all experimental steps involved. This model type could thus be a viable option for breeding programs and their collaborators who currently conduct static digestion methods. The PS is also more readily scalable across institutions than the HGS; computer-aided design specifications for the PS are available in Swackhamer et al..^9^ Further optimizing the dynamic features of the PS to enable real-time inputs and outputs would facilitate the emulation of gastric juice secretion (an input), gastric emptying (an output), and collection of samples from a given tube (an output) at intermediate time points throughout digestion. Real-time nutrient sensors ^89^ in the digestion apparatus could also be advantageous in multiple of the digestion methods tested herein; e.g., for continuous (and non-destructive) monitoring of trait values.

### 4.4 Operational Considerations and Recommendations for use of Dynamic Digestion Models in Plant Breeding

The availability of only small sample masses from small-plot field research trials is a common feature in breeding programs, particularly from previous growing seasons due to limits on physical storage space. As such, a dynamic digestive platform that requires even smaller sample masses than the simplified HGS method (i.e., <100 grams, but still enough to be representative of each plot) could be helpful for the adoption of these platforms. The sample mass for the full HGS is designed to represent a full meal (or serving size for human consumption). Models using substantially smaller sample masses (such as the PS) will need to be routinely monitored for emulation of both physical/mechanical and chemical processes.

Additionally, the samples used for aim 2 were harvested in 2019 and had been stored in room-temperature conditions (a standard practice for grain samples in breeding programs due to sample volume), which could have impacted phenolic levels.^90^ For UC Southwest Gold raw beans, total phenolics were comparable in samples from San Gregorio in 2022 vs. Pescadero in 2019. When comparing digestive phenolics from aims 1 and 2 (Figures 5 and 7), particularly at the 240-minute time point for the simplified HGS method, the beans from 2022 used in aim 1 showed more phenolics than those from 2019 used for aim 2. Long-term storage under non-ideal conditions can cause quality defects such as hard-shell and hard-to-cook characteristics, which may lead to decreased digestibility and nutrient bioavailability.^91^ However, food scientific examinations have often purchased samples from commercial sources (e.g., retail stores) with unknown harvest dates (and unknown genotypes and growing environments, in some cases). We would recommend conducting simulated gastrointestinal digestion on freshly harvested dry beans of known genotypes and production environments, when feasible, to control these experimental factors.

Finally, the analyses used to assess nutrient values for this study in pre-digestion samples and samples extracted from digestions were expensive and time-consuming. Using near-infrared spectroscopy to predict at-harvest nutrient values in larger sample sets could be helpful in selecting a subset of samples for dynamic digestion, if it is not possible to digest all samples of interest via a higher-throughput dynamic method such as the PS. It would also be informative to test whether high-throughput *in vitro* protein digestibility assays can partially predict levels of free amino groups in dynamic digestive models; if so, such assays could also be used in selection of samples for dynamic digestion. For example, the rapid protocol published by Diatta-Holgate et al. for analysis of cooked sorghum or maize used a culturally relevant preparation method (porridge from flour) and a two-hour incubation with pepsin, with throughput of 280 samples in two days.^92^ Alongside breeding for direct consumption of whole beans, samples with favorable nutrient profiles such as high protein or starch at harvest (e.g., in ground flour), high phenolic concentrations in the broth, or favorable patterns of protein and starch hydrolysis during digestion could have utility as ingredients. This area of research and product development would benefit from sustained collaborations between plant breeders and food scientists.^93^

Testing of samples from more genotypes and market classes on dynamic digestion platforms would further inform breeding efforts. In such studies, we would recommend incorporating an empirical evaluation of cooking time ^94–97^ and continued use of genotype-specific cooking times to maximize comparability of digestive outcomes across samples; the effects of environment on cooking time have been found to be less major.^95,98^ These dynamic digestion platforms could also inform research and development efforts in other grain legumes. While puree and crackers produced from a commercial supply of garbanzos (*Cicer arietinum* L.) have been tested on the HGS,^6^ examination of diverse and elite cultivars of various grain legume species from diverse growing environments would be a major next step. Testing a larger number of samples (representing genotype-environment combinations) would also be helpful to determine whether rank-order differences in nutrient release are observed across multiple digestion platforms. That is, if a higher-throughput platform incurs less release of one or more nutrients but the resulting trait values are highly correlated with those from the HGS, the former could still have utility in breeding (as the same ‘best’ genotypes would be identified, which could be used as varietal candidates and/or parents in breeding pipelines).

### 4.5 Conclusion

In summary, dynamic simulated digestion methods have herein provided quantitative information on bioaccessible nutrient profiles and their evolution during digestion in common bean, an important protein staple for direct consumption by humans. The understanding gained through this work, and the methods described herein, will assist breeders in their efforts to improve the bioaccessible nutrient profiles that are offered to consumers of these crops.

## Conflicts of Interest

There are no conflicts of interest to declare.

## Acknowledgments

We gratefully acknowledge Troy Williams for development of the ImageJ macro, Clay Swackhamer for training on the Peristaltic Simulator, and growers (including R.C.) for hosting field trials.

## Abbreviations

DNS: 3,5-Dinitrosalicylic Acid
DW: Dry Weight
FC: Folin-Ciocâlteu
FRAP: Ferric Reducing Antioxidant Power
GJ: Gastric Juice
HGS: Human Gastric Simulator
IF: Intestinal Fluid
NA: Not Applicable
OPA: o-Phthaldialdehyde
PS: Peristaltic Simulator
RDS: Reducing Sugars
SD: Standard Deviation
UC: University of California

## Notes

### Competing Interest Statement

The authors have declared no competing interest.

