## Supplemental Tables and Figures for "An empirical evaluation of simulated gastrointestinal digestion platforms for use in plant breeding, using common bean (*Phaseolus vulgaris* L.) as a model"

**Supporting Information description:** The Supporting Information includes supplemental figures (7), supplemental tables (3), and a supplemental file (1).

**Supporting Information:**

- Supplemental File S1. Results of mixed ANOVA and Tukey's Honestly Significant Difference post hoc tests for each trait. Uploaded as a separate file due to file size.

Table S1: Sources for all chemical reagents used.

| Chemical Name | Chemical Formula | Source |
|---|---|---|
| Sodium Chloride | $NaCl$ | Fisher Chemical CAS: 7647-14-5 |
| Methanol | $CH_3OH$ | Fisher Chemical CAS: 67-56-1 |
| Sodium Carbonate | $Na_2CO_3$ | Fisher Chemical CAS: 497-19-8 |
| Folin-Ciocâlteu | $C_{10}H_5NaO_5S$ | Sigma-Aldrich CAS: 12111-13-6 |
| Gallic Acid | $C_7H_6O_5$ | Acros Organics CAS: 149-91-7 |
| TPTZ | $C_{18}H_{12}N_6$ | Chem-Impex Int'l Inc. CAS: 3682-35-7 |
| Hydrochloric Acid | HCL | Sigma-Aldrich CAS: 7647-01-0 |
| iron(3+) trichloride hexahydrate | $FeCl_3(H_2O)_6$ | Fisher Science Education CAS: 10025-77-1 |
| Sodium Acetate Trihydrate | $CH_3COONa(H_2O)_3$ | Amresco CAS: 6131-90-4 |
| Acetic Acid pH 3.6 | $CH_3COOH$ | Spectrum Chemical MFG. Corp. CAS: 64-19-7 |
| Trolox | $C_{14}H_{18}O_4$ | Sigma-Aldrich CAS: 53188-07-1 |
| 3,5-Dinitrosalicylic acid | $C_7H_4N_2O_7$ | Sigma Life Science CAS: 609-99-4 |
| Sodium potassium tartrate | $KNaC_4H_4O_6(H2O)_4$ | Fisher Chemical CAS: 6381-59-5 |
| Sodium hydroxide | $NaOH$ | Fisher Chemical CAS: 1310-73-2 |
| Maltose | $C_{12}H_{22}O_{11}$ | Alfa Aesar CAS: 6363-53-7 |
| Sodium dodecyl sulfate | $NaC_{12}H_{25}SO_4$ | Sigma-Aldrich CAS: 151-21-3 |

| | | |
|---|---|---|
| Sodium tetraborate decahydrate | $Na_2B_4O_7(H_2O)_{10}$ | Sigma-Aldrich CAS: 1303-96-4 |
| *O*-phthaldialdehyde (OPA) | $C_6H_4(CHO)_2$ | Sigma-Aldrich CAS: 643-79-8 |
| 2-mercaptoethanol | $HOCH_2CH_2SH$ | Bio-Rad CAS: 60-24-2 |
| Glycine | $C_2H_5NO_2$ | Bio-Rad CAS: 56-40-6 |

Table S2: Composition of fluids for simulated digestions.

| Component | Saliva | Gastric Juice | Intestinal Fluid |
|---|---|---|---|
| pH | 7.0 | 0.8 | 7.0 |
| Enzyme | α-amylase<br>mpbio CAS: 9000-90-2 | Pepsin<br>mpbio CAS: 9001-75-6 | Pancreatin<br>Sigma-Aldrich CAS: 8049-47-6 |
| | 1.18 mg/mL | 0.008 mg/mL | 2.4 mg/mL |
| Mucin<br>Sigma-Aldrich CAS: 84082-64-4 | 1.0 mg/mL | 1.5 mg/mL | NA |
| Bile Extract<br>Sigma-Aldrich CAS: 8008-63-7 | NA | NA | 10 mg/mL |
| $CaCl_2(H_2O)_2$<br>Fisher BioReagents CAS: 10035-04-8 | 0.22 mg/mL | 0.022 mg/mL | 0.088 mg/mL |
| KCl<br>Fisher Scientific CAS: 7447-40-7 | 1.13 mg/mL | 0.515 mg/mL | 0.507 mg/mL |
| $KH_2PO_4$<br>Fisher Scientific CAS: 7778-77-0 | 0.5 mg/mL | 0.122 mg/mL | 0.109 mg/mL |
| $NaHCO_3$<br>Ward's Science CAS: 144-55-8 | 1.14 mg/mL | 2.1 mg/mL | 7.14 mg/mL |
| NaCl<br>Fisher Chemical CAS: 7647-14-5 | NA | 2.761 mg/mL | 2.25 mg/mL |
| $MgCl_2(H_2O)_6$<br>Fisher Scientific CAS: 7791-18-6 | 0.03 mg/mL | 0.024 mg/mL | 0.067 mg/mL |
| $(NH_4)_2CO_3$<br>Acros Organics CAS: 506-87-6 | 0.01 mg/mL | 0.048 mg/mL | NA |

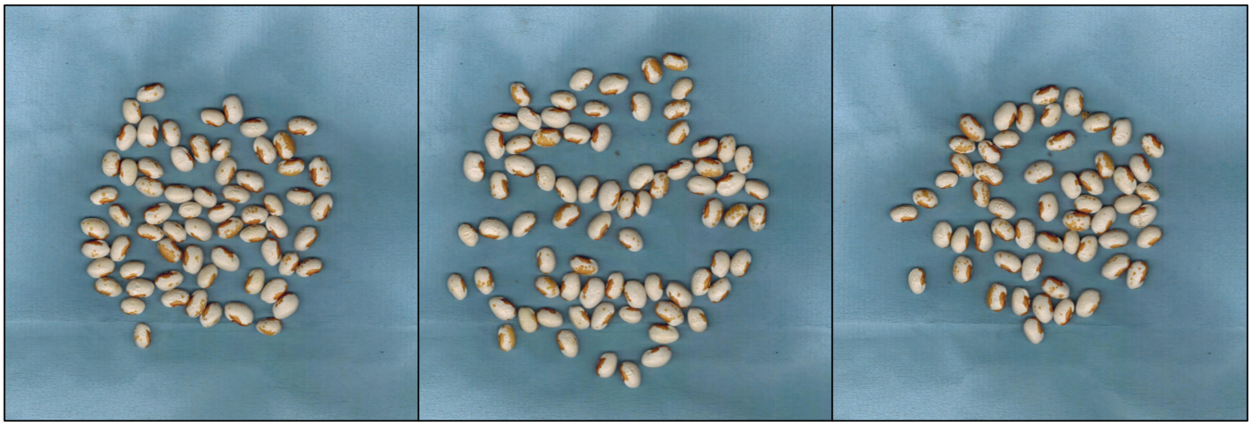

Figure S1. Seed from UC Southwest Gold produced in San Gregorio in 2022, as depicted in the by-plot (i.e., for each biological replicate) images used for seed coat pigmentation analysis via ImageJ macro.

Table S3. Raw data for seed coat coloration that was produced by the ImageJ macro. The traits reported are the percentage of pigmented area in the scan, the percentage of the image that is seed, and the proportion of the seed that is pigmented (which is calculated by dividing the first trait by the second trait). The latter trait, multiplied by 100, is reported in the text as the percentage of pigmented seed coat area. Bold text represents average values and standard deviation for each sample type.

| Genotype | Plot | Location | Year | Pigment % Area in Scan | % of Image that is Seed | Proportion of Seed Pigmented |
|---|---|---|---|---|---|---|
| UC Southwest Gold | 1 | Blue House Farm (San Gregorio, CA) | 2022 | 1.302 | 6.973 | 0.187 |
| UC Southwest Gold | 2 | Blue House Farm (San Gregorio, CA) | 2022 | 1.569 | 9.401 | 0.167 |
| UC Southwest Gold | 3 | Blue House Farm (San Gregorio, CA) | 2022 | 1.409 | 7.952 | 0.177 |
| | | | | **1.427 ± 0.134** | **8.109 ± 1.222** | **0.177 ± 0.010** |
| UC Southwest Gold | 1 | Fifth Crow Farm (Pescadero, CA) | 2019 | 1.194 | 11.593 | 0.103 |
| UC Southwest Gold | 2 | Fifth Crow Farm (Pescadero, CA) | 2019 | 1.215 | 13.052 | 0.093 |
| UC Southwest Gold | 3 | Fifth Crow Farm (Pescadero, CA) | 2019 | 0.982 | 11.546 | 0.085 |
| | | | | **1.130 ± 0.129** | **12.064 ± 0.856** | **0.094 ± 0.009** |
| UC Southwest Gold | 1 | Student Farm (Davis, CA) | 2019 | 8.526 | 13.834 | 0.616 |
| UC Southwest Gold | 2 | Student Farm (Davis, CA) | 2019 | 7.028 | 10.843 | 0.648 |
| UC Southwest Gold | 3 | Student Farm (Davis, CA) | 2019 | 15.983 | 24.758 | 0.646 |
| | | | | **10.512 ± 4.797** | **16.478 ± 7.325** | **0.637 ± 0.018** |
| UC Southwest Red | 1 | Fifth Crow Farm (Pescadero, CA) | 2019 | 3.675 | 13.815 | 0.266 |
| UC Southwest Red | 2 | Fifth Crow Farm (Pescadero, CA) | 2019 | 1.377 | 9.075 | 0.152 |
| UC Southwest Red | 3 | Fifth Crow Farm (Pescadero, CA) | 2019 | 2.778 | 13.000 | 0.214 |
| | | | | **2.610 ± 1.158** | **11.963 ± 2.534** | **0.211 ± 0.057** |
| UC Southwest Red | 1 | Student Farm (Davis, CA) | 2019 | 13.145 | 18.598 | 0.707 |
| UC Southwest Red | 2 | Student Farm (Davis, CA) | 2019 | 13.240 | 18.475 | 0.717 |
| UC Southwest Red | 3 | Student Farm (Davis, CA) | 2019 | 17.909 | 25.549 | 0.701 |
| | | | | **14.765 ± 2.723** | **20.874 ± 4.049** | **0.708 ± 0.008** |

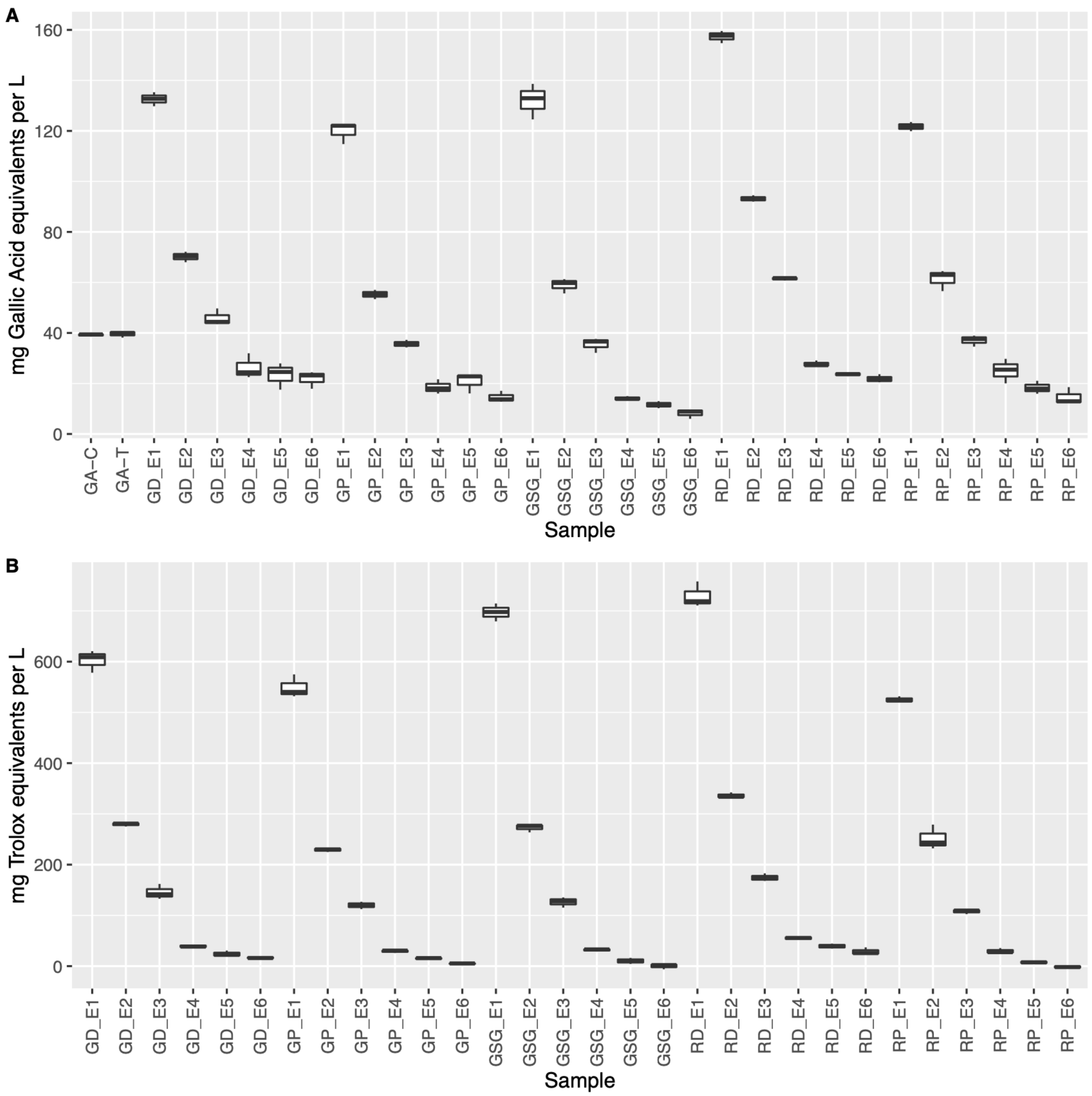


Figure S2. A) Phenolic extraction fractions analyzed via the FC assay for quantification of total phenolics in raw bean samples. B) Antioxidant power fractions analyzed via the FRAP assay for quantification of total antioxidant power in raw bean samples. GA-C represents a gallic acid control sample, while GA-T represents a gallic acid test sample which underwent the treatment of all six rounds of extraction. Samples with a first letter of 'G' are of the UC Southwest Gold variety. Samples with a first letter of 'R' are of the UC Southwest Red variety. Samples with a second letter of 'D' were grown in Davis, while 'P' denotes samples grown in Pescadero and 'SG" denotes samples grown in San Gregorio. The E1-6 suffix represents subsequent extraction fractions for a given sample, where E1 represents the first round of extraction and E6 represents the sixth (final) round of extraction.

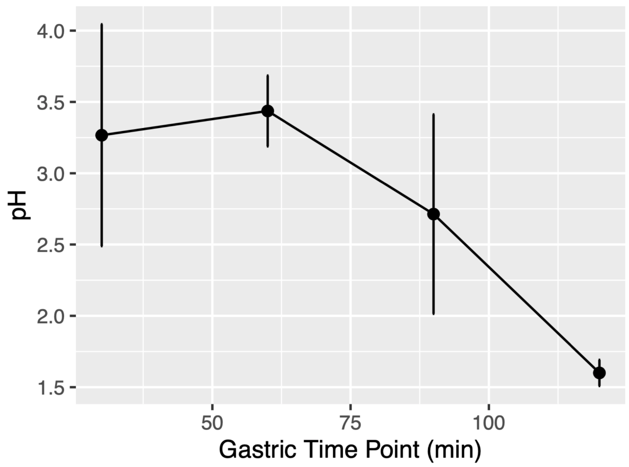

Figure S3: Full HGS gastric emptying pH curve.

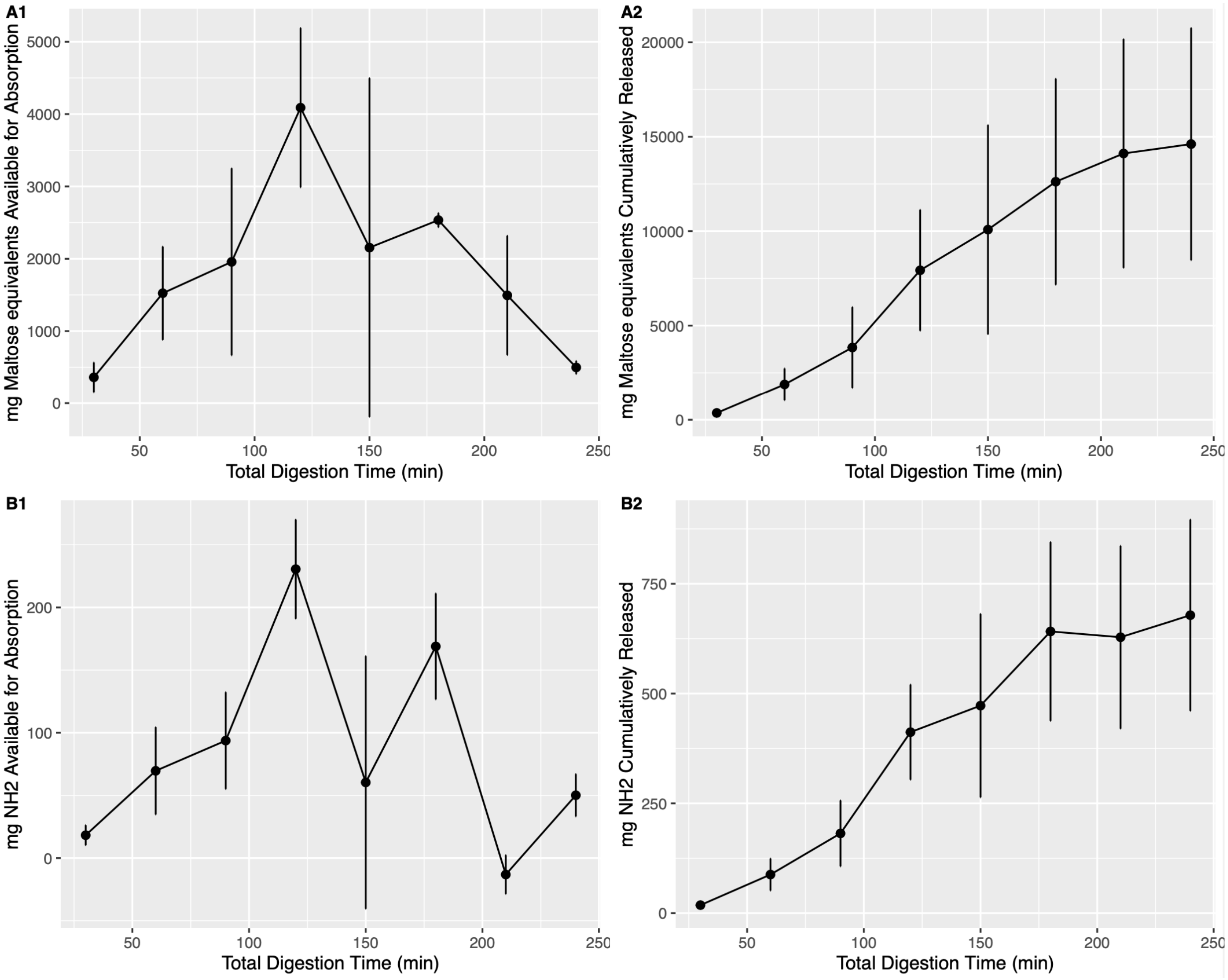

Figure S4: Full HGS absorption curves (A1 and B1 depict values at each 30-minute time point; A2 and B2 depict cumulative values) for starch and protein hydrolysis as determined by RDS and OPA assays, respectively.

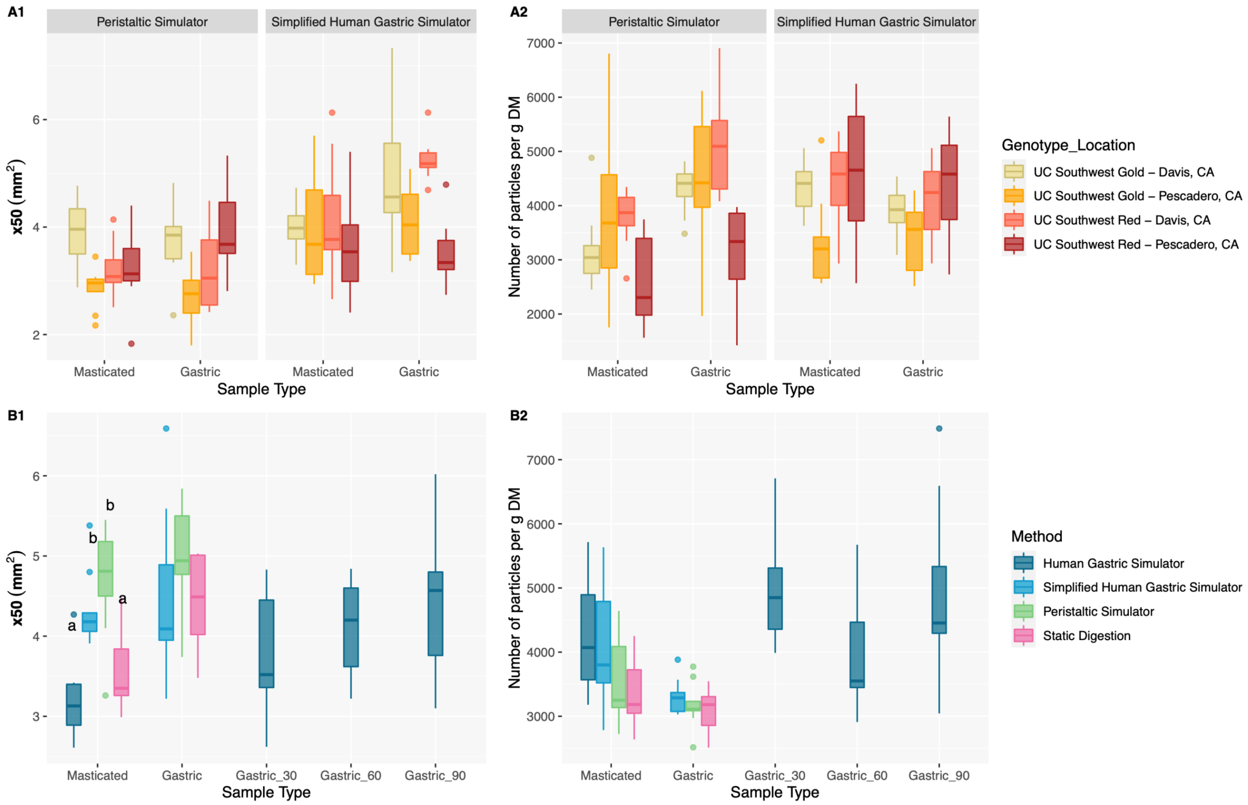


Figure S5: Particle size data for masticated and gastric samples from A) the UC Southwest variety comparison using both the simplified HGS and PS digestion methods, and B) the four-way simulated digestion method comparison. Panel 1 depicts the size of a hypothetical sieve through which 50% of the particle area could pass (x50). Panel 2 depicts the calculated number of particles per gram dry weight of sample. 'Gastric' denotes the 120-minute time point at the end of the gastric phase for the simplified HGS, PS, and static methods, while the additional 30-, 60-, and 90-minute gastric time points (with numbers appearing as suffixes along the X-axis) represent the full HGS method time points for technical replicates from which there was enough sample available for collection. Significance was determined using a Tukey's Honestly Significant Difference post hoc test following mixed ANOVA for comparable groups. Letters were not displayed for time points within which no significant differences were observed. No significant differences were observed in any of the pairwise comparisons of group means in panels A1, A2, and B2, such that connecting letters are not depicted in these panels.

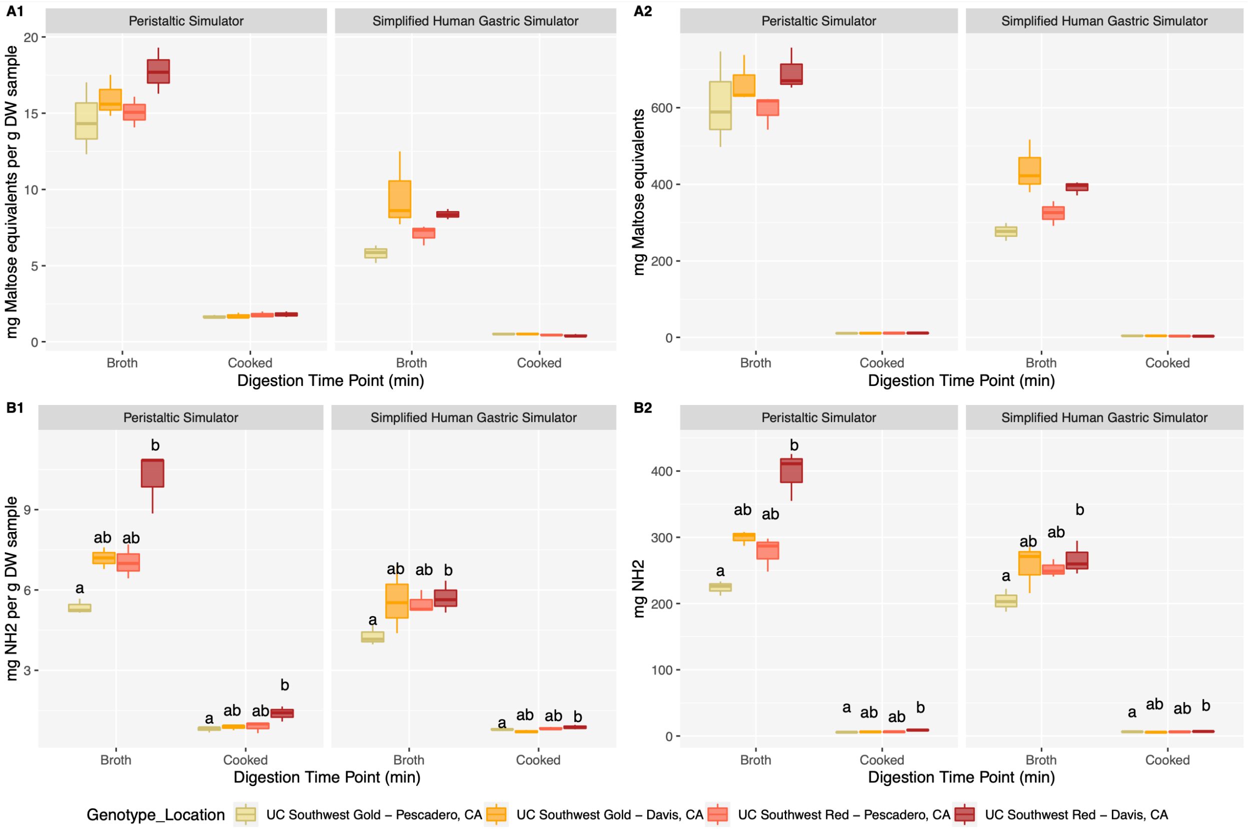


Figure S6. Starch and protein hydrolysis in the broth and cooked bean samples for the UC Southwest Red and Gold variety comparison using both the simplified HGS and PS digestion methods. A) Starch hydrolysis was determined via RDS assays. B) Protein hydrolysis was determined via OPA assays. Significance was determined using a Tukey's Honestly Significant Difference post hoc test following mixed ANOVA. No significant differences were observed in any of the pairwise comparisons of group means in panels A1 and A2, such that connecting letters are not depicted in these panels.

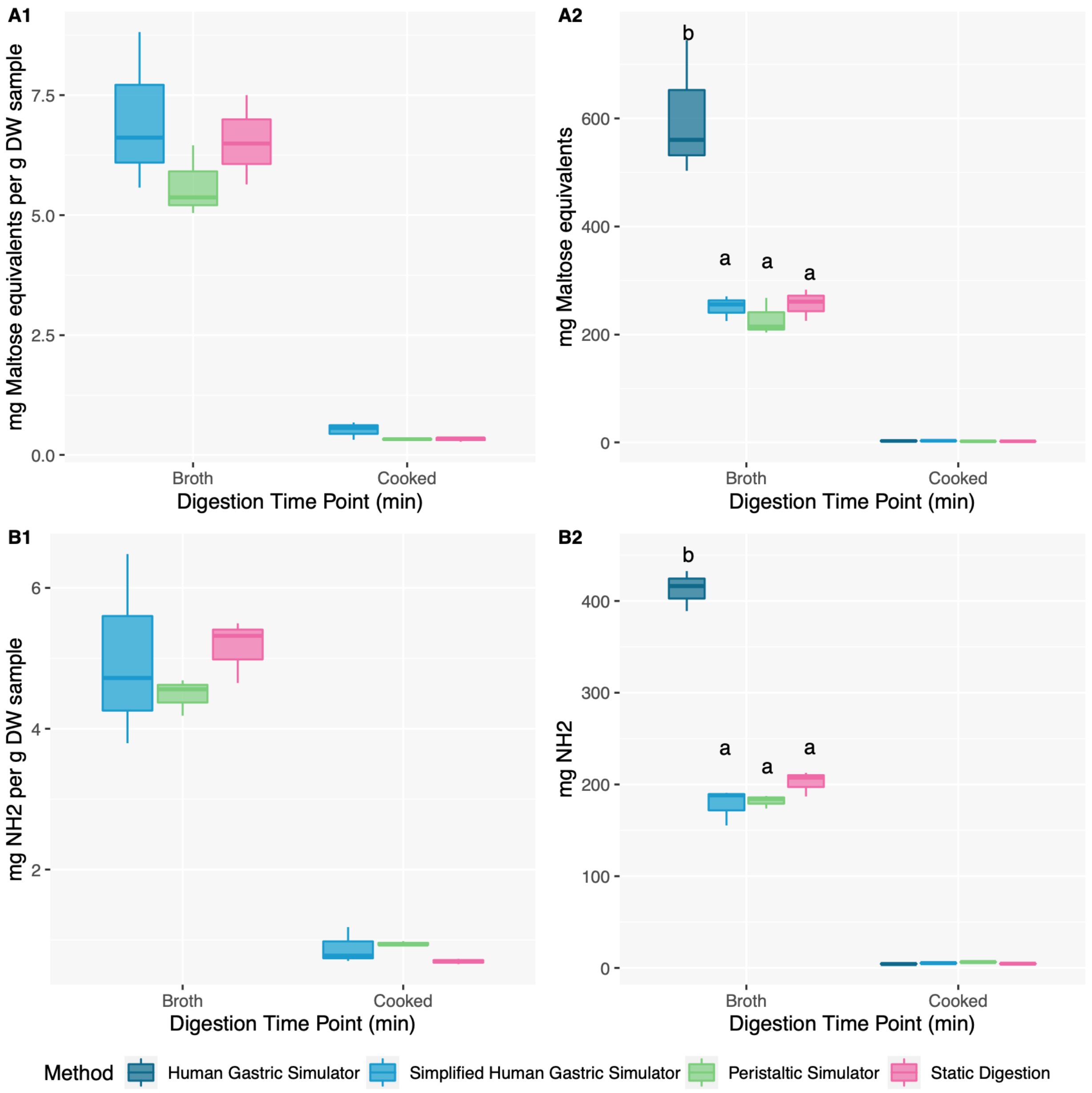


Figure S7. Starch and protein hydrolysis in the broth and cooked bean samples for the four-way simulated digestion method comparison. A) Starch hydrolysis was determined via RDS assays. B) Protein hydrolysis was determined via OPA assays. Significance was determined using a Tukey's Honestly Significant Difference post hoc test following mixed ANOVA. Comparisons were made within time points, and letters were not displayed for time points within which no significant differences were observed. No significant differences were observed in any of the pairwise comparisons of group means in panels A1 and B1, such that connecting letters are not depicted in these panels.

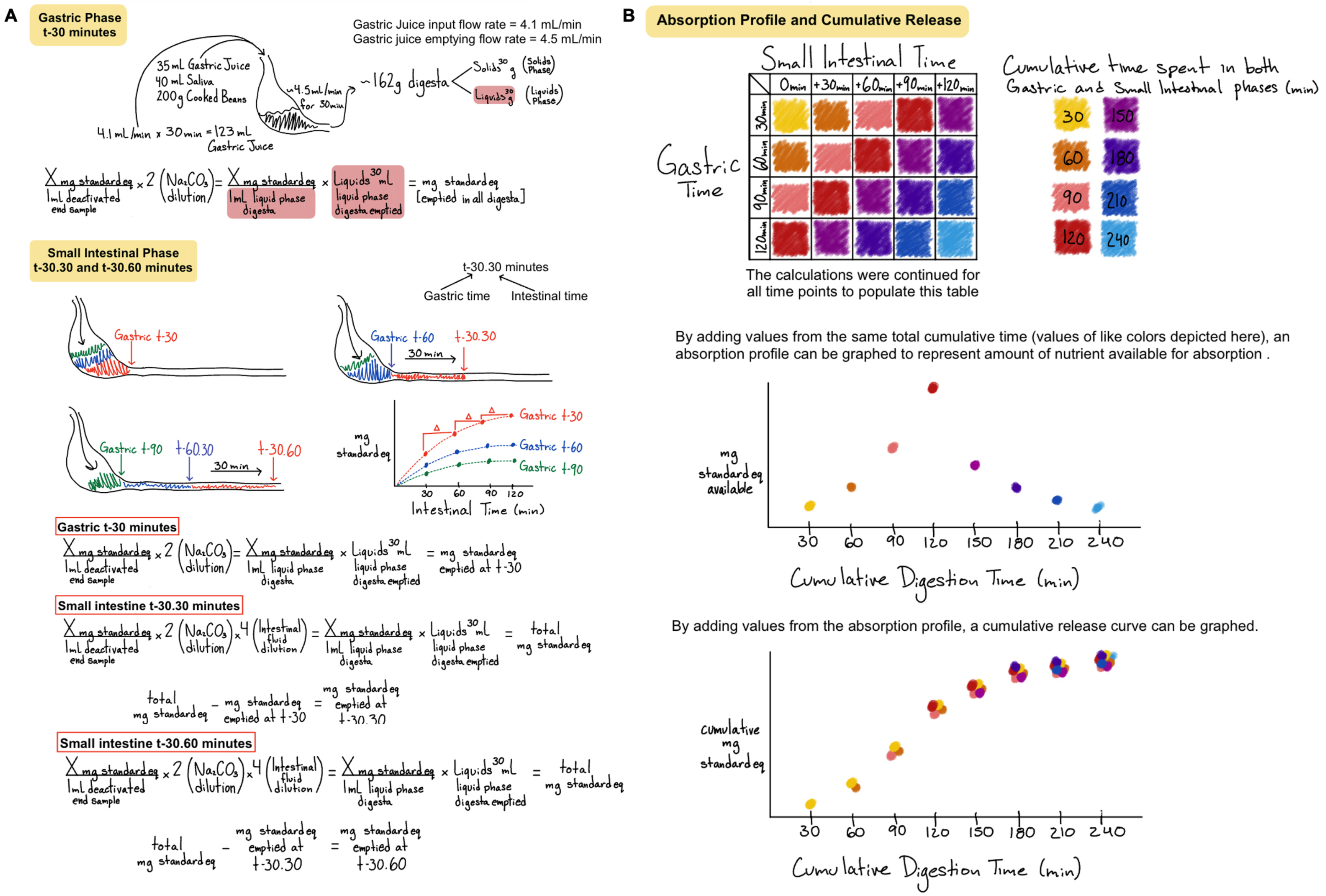

Figure S8. Calculations used to determine analyte concentrations throughout the full HGS method.
